# Functional diversification of hybridoma produced antibodies by CRISPR/HDR genomic engineering

**DOI:** 10.1101/551382

**Authors:** Johan M.S. van der Schoot, Felix L. Fennemann, Michael Valente, Yusuf Dolen, Iris M. Hagemans, Anouk M.D. Becker, Camille M. Le Gall, Duco van Dalen, Alper Cevirgel, J. Armando C. van Bruggen, M Engelfriet, Tomislav Caval, Arthur E.H. Bentlage, Marieke F. Fransen, Maaike Nederend, Jeanette H.W. Leusen, Albert J.R. Heck, Gestur Vidarsson, Carl G. Figdor, Martijn Verdoes, Ferenc A. Scheeren

**Affiliations:** Department of Tumor Immunology, Radboud Institute for Molecular Life Sciences, Radboud University Medical Center, Geert Grooteplein 26, 6525 GA Nijmegen, The Netherlands; Division of Immunology, The Netherlands Cancer Institute, Plesmanlaan 121, 1066 CX, Amsterdam, The Netherlands; Biomolecular Mass Spectrometry and Proteomics, Bijvoet Center for Biomolecular Research and Utrecht Institute for Pharmaceutical Sciences, Utrecht University, Padualaan 8, 3584 CH Utrecht, The Netherlands; Sanquin Research, Department of Experimental Immunohematology, Amsterdam, The Netherlands, and Landsteiner Laboratory, Amsterdam UMC, University of Amsterdam, Amsterdam, Plesmanlaan 125, Amsterdam 1066 CX, The Netherlands; Department of Immunohematology and Blood Transfusion, Leiden University Medical Center (LUMC), Albinusdreef 2, 2333 ZA Leiden, The Netherlands; Laboratory for Translational Immunology, UMC Utrecht, Utrecht, The Netherlands; Department of Medical Oncology, Department of Surgery and Department of Immunohematology and Blood Transfusion, Leiden University Medical Center (LUMC), Albinusdreef 2, 2333 ZA Leiden, The Netherlands

## Abstract

Hybridoma technology is instrumental for the development of novel antibody therapeutics and diagnostics. Recent preclinical and clinical studies highlight the importance of antibody isotype for therapeutic efficacy. However, since the sequence encoding the constant domains is fixed, tuning antibody function in hybridomas has been restricted. Here, we demonstrate a versatile CRISPR/HDR platform to rapidly engineer the constant immunoglobulin domains to obtain recombinant hybridomas which secrete antibodies in the preferred format, species and isotype. Using this platform, we obtained recombinant hybridomas secreting Fab’ fragments, isotype switched chimeric antibodies, and Fc-silent mutants. These antibody products are stable, retain their antigen specificity, and display their intrinsic Fc-effector functions in vitro and in vivo. Furthermore, we can site-specifically attach cargo to these antibody products via chemo-enzymatic modification. We believe this versatile platform facilitates antibody engineering for the entire scientific community, empowering preclinical antibody research.

**One Sentence Summary:** We demonstrate a universal CRISPR/HDR based platform for rapid genetic engineering of hybridomas to obtain functionally diverse antibody isotype panels in the species and format of choice.

## Introduction

Monoclonal antibodies (mAbs) have revolutionized the medical field, allowing the treatment of diseases, which were previously deemed incurable (*1*). For mAb discovery, screening and production, hybridoma technology has been widely used since its introduction in 1975 (*2*). Hybridomas are immortal cell lines capable of secreting large quantities of mAbs and have been proven to be essential for the development of new antibody-based therapies. Over the past decades, a large number of hybridomas have been generated, validated and made available, including hybridomas used for preclinical research. Besides the antigen specificity, the mAb format and isotype are important determinants for performance in preclinical models. However, as the mAb heavy chain sequence is genetically fixed in the hybridoma, the possibilities to adjust antibody format and functionality without compromising target specificity are limited. Typically, genetically engineered mAbs are produced using recombinant technology. This requires sequencing of the variable domains, cloning of these variable domain sequences into the plasmids with the appropriate backbone, and expression of these plasmids in transient systems. These procedures are often time-consuming, challenging and expensive and therefore often outsourced to specialized contract research companies, hampering academic early-stage antibody development and preclinical application.

The importance of the antigen binding variable regions is apparent. However, the constant domains which form the fragment crystallisable (Fc) domain are also central to the therapeutic efficacy of mAbs. Numerous studies have demonstrated that therapeutic efficacy of certain mAbs depends on the Fc domain and their engagement with specific Fc receptors (FcRs) (*3*–*8*). In a recent example, Vargas et al. demonstrated that anti-CTLA-4 immunotherapeutic working mechanism revolves around depletion of regulatory T-cells in the tumor microenvironment, utilizing Fc variants of the same antibody (*4*). The authors postulated that this could explain why cancer patients treated with ipilimumab, a human IgG1 CTLA-4 targeting mAb, greatly benefit from an activating FcγRIIIa-polymorphism which increases affinity for human IgG1 (*4*). This highlights the central role the Fc plays in antibody-based therapeutics, and emphasizes the importance of evaluating novel mAbs in different Fc formats in early development.

The clustered regularly interspaced short palindromic repeat (CRISPR)/Cas9 (CRISPR-associated Protein 09) targeted genome editing technology opened up a plethora of exciting opportunities for gene therapy, immunotherapy, and bioengineering (*9*). Recently, CRISPR-Cas9 has been used to modulate mAb expression in hybridomas: Cheong *et al*. forced class-switch recombination in IgM producing hybridomas to downstream immunoglobulin classes via deletion of the natural S regions, and induced Fab’ fragment production by deletion of the entire Fc region (*10*). Pogson *et al*. generated a hybridoma platform in which the variable domains can be substituted (*11*), and affinity maturation can be simulated via homology directed mutagenesis with single stranded oligos (*12*). Khoshnejad *et al*. engineered a hybridoma to introduce a sortase recognition motif (sortag) (*13*) on the C-terminus of the constant heavy chain domain 3 (CH3) for site-specific chemoenzymatic antibody modification purposes (*14*). However, to date no platform has been described for versatile and effective Fc substitution in hybridomas with constant domains from foreign species, isotypes or formats.

Here, we demonstrate genetic engineering of hybridomas via a one-step CRISPR/ Homology Directed Repair (HDR)-based strategy enabling rapid generation of recombinant hybridomas secreting mAbs in the Fc format of choice with the freedom to install preferred tags or mutations. We generated hybridomas producing Fab’ fragments, c-terminally equipped with a sortag and a histag for site-specific chemoenzymatic modification and purification purposes. Furthermore, we incorporated novel constant domains upstream of the native constant region to generate hybridomas which secrete chimeric antibodies with the isotype and species of choice, as well as Fc mutant mAbs without compromising antigen specificity. The engineered antibodies are easily isolated from the hybridoma supernatant and display their expected biochemical and immunological characteristics, both in vitro and in vivo. Because we target the constant domains of the immunoglobulin locus, our CRISPR/HDR method is adaptable to hybridomas from any species or isotype. We believe that this versatile platform will empower preclinical antibody research by opening up antibody engineering for the entire scientific community.

## Results

### Generation of recombinant hybridomas producing sortagable Fab’ fragments

Monovalent Fab’ fragments consist of a shortened heavy chain and light chain and, due to their smaller size, have certain advantages over their parental mAbs. Fab’ fragments have better tissue penetration, are not susceptible to Fc-mediated uptake and do not exert immune effector functions. Therefore, Fab’ fragments have been used for treatment of certain autoimmune diseases (*15*), targeting of different payloads (*16, 17*), and generation of bispecific antibodies (*18*). The conventional method to obtain Fab’ fragments is papain cleavage of the parental mAb followed by purification of the Fab’ fragment. However, this does not allow for installment of useful tags or mutations. Alternatively, one could use recombinant production in transient systems, however this requires prior knowledge of the variable heavy and light chain sequence, optimization of the transient system, and multiple rounds of production. Rapid conversion of hybridomas into Fab’ secreting cell lines via CRISPR/HDR engineering represents a simple and cost-effective alternative to obtain inexhaustible source of Fab’ fragment with the freedom, to introduce mutations or tags of choice.

To obtain Fab’ secreting hybridomas via CRISPR/HDR we selected the hybridoma clone NLDC-145 (*19*) as a first target (**Fig. 1A,B**). This hybridoma secretes mAbs of the rat IgG2a (rIgG2a) isotype, and is specific for murine DEC205. This endocytic receptor is abundantly expressed by dendritic cells, and employed as a target in various vaccination strategies (*20, 21*). To target the heavy chain locus of NLDC-145, we used an optimized guide RNA (gRNA-H) which directs Cas9 to the rIgG2a hinge region to generate double stranded breaks (**Fig. S1**). To repair the double stranded break, we designed an HDR donor construct (**Table S1**) that inserts a sortag and histag directly upstream of the cysteine involved in heavy chain dimer formation. Furthermore, we included a *blasticidin-S deaminase* (*Bsr*) gene to select cells, which successfully integrated the construct. We electroporated NLDC-145 hybridoma cells with the HDR Fab’ donor construct and Cas9 PX459 vector containing gRNA-H, and subsequently applied blasticidin pressure. After blasticidin selection, we assessed knock-in of the donor construct by genomic PCR with a forward primer that hybridizes upstream of the 5’ homologous arm, and a reverse primer specific for the HDR construct. PCR products of the expected size (∼1600bp) were exclusively observed for the transfected population, indicating integration of the donor construct in the correct genomic location (**Fig. 1C**). After one week, we performed a limiting dilution to obtain monoclonal cell suspensions. We used the supernatant of these monoclonal cell suspensions to determine antigen specificity and phenotype of the secreted product. We first stained JAWSII, a DEC205 expressing cell line, with the supernatant from each clone and performed a subsequent secondary staining for the histag and rIgG2a heavy chain. Flow cytometry analysis revealed that a large fraction of monoclonal hybridomas was successfully engineered (**Fig. 1D**), as we observed DEC205 binding constructs harboring a histag in the supernatant of 47 out of 76 clones. In contrast, only a single cell suspension (clone 48) still produced the mAb DEC205 of the rat IgG2a isotype. To validate the flow cytometry results, we performed a Western Blot for the heavy and light chain on the supernatant of clones 47 to 52 (**Fig. 1E**). The four clones that were histag^pos^ rIgG2a^neg^ by flow cytometric analysis, indeed secreted a shortened heavy chain that was histag positive, confirming production of histagged Fab’ fragments by these clones. Clone 48 (histag^neg^, rIgG2a^pos^) retained its expression of the rat heavy chain, while for clone 47 (histag^neg^, rIgG2a^neg^) only rat light chain could be observed.

**Figure 1.**
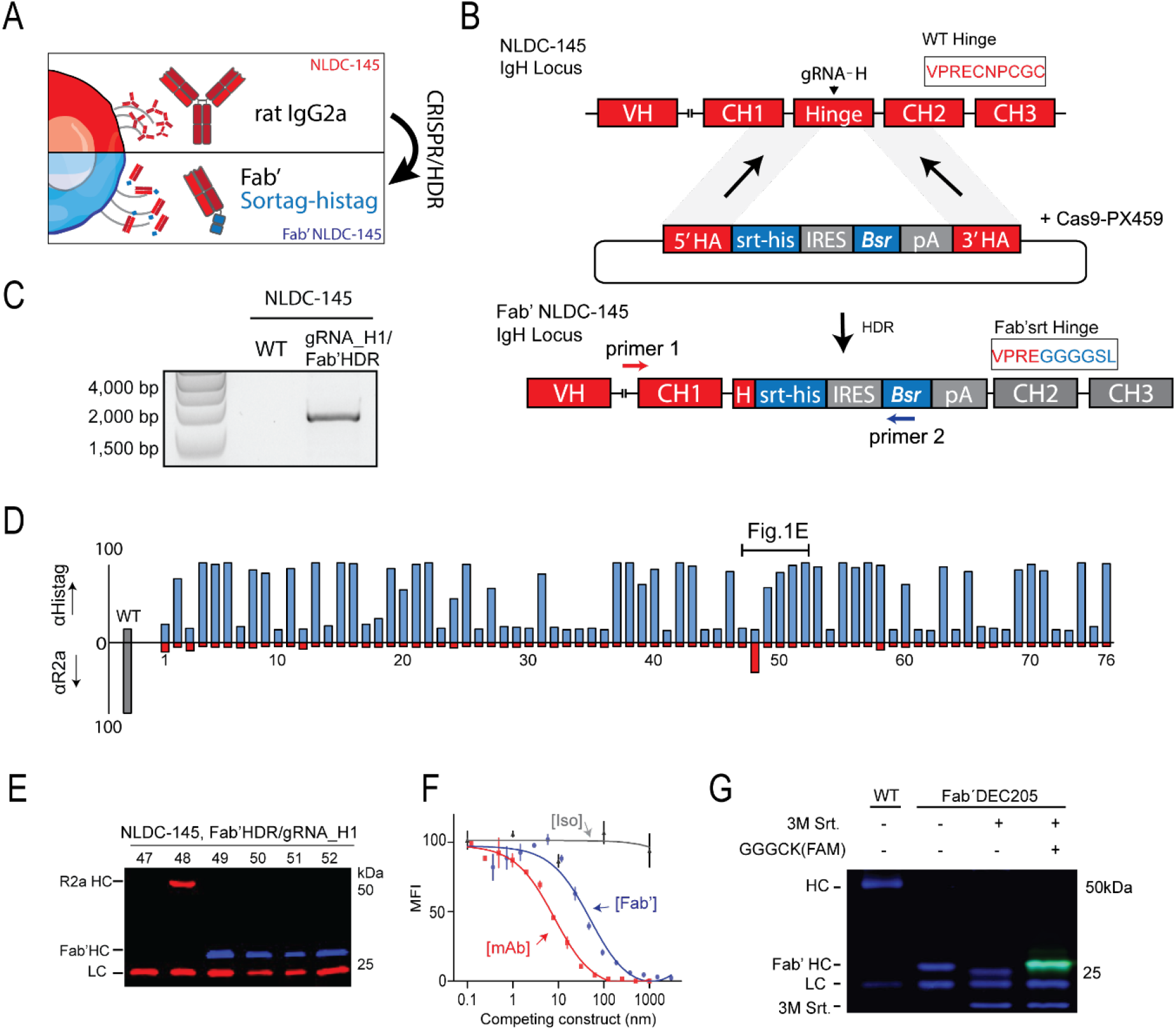
CRISPR/HDR engineering of hybridoma NLDC145 to obtain sortagable Fab’ fragments against DEC205. **(A)** Schematic representation of the CRISPR/HDR approach to convert wildtype hybridomas to Fab’ fragment producing cell lines. **(B)** Shown is the targeted IgH locus of NLDC-145 with the variable region (VH) and constant regions (CH1, Hinge, CH2, CH3,) annotated. Cas9 is guided by gRNA-H to hinge region and creates a double stranded break before the first cysteine. The double stranded break is subsequently repaired by homology directed repair via the donor construct, consisting of; a sortag and histag motif (srt-his), an internal ribosomal entry site (IRES), *blasticidin-S deaminase* (*Bsr*), polyA transcription termination signal (pA) and homology arms (5’ HA, 3’ HA). The first 10 amino acids of the hinge, before and after successful CRISPR/HDR are indicated. **(C)** Three days after electroporation PCR was performed on genomic DNA from WT and CRISPR/HDR targeted population using primers 1 and 2 (Fig. 1b). Agarose gel shows amplicon of expected size exclusively for the CRISPR/HDR targeted population, indicating correct integration of the donor construct within the population. **(D)** After limiting dilution of CRISPR-HDR targeted cells, a flow cytometry screen was performed on the supernatant of monoclonal cell suspensions. To this end, DEC-205 expressing cells (JAWSII) were incubated with clonal supernatants in combination with secondary antibodies against histag (blue) or rat IgG2a (red). Exclusive histag signal indicates production of Fab’ fragments. **(E)** Immunoblotting of the supernatant of clone 47-52 for histag (blue) and rat heavy and light chain (red) confirms flow cytometry screen. **(F)** Competition assay, JAWSII were incubated with a serial dilution (150µg/mL-20ng/mL) of either purified Fab’DEC205-srt (blue), αDEC205 mAb (red), or an isotype control (grey) in combination with 1µg/mL of fluorescently labelled αDEC205 mAb. Decrease in mean fluorescent intensity relates to increase in competition for DEC-205 binding with fluorescent labelled αDEC205 mAb. *n*=3, mean ± s.e.m **(G)** Fab‘DEC205-srt can be c-terminally functionalized with a fluorescently labelled probe (GGGCK(FAM)) by using sortase (3m Srt.) mediated ligation.

For further characterization of the secreted Fab’DEC205-sortag-histag (hereafter Fab’DEC205srt) we selected one clone (hereafter Fab’ hybridoma) of which the heavy chain sequence was validated by Sanger sequencing. After expansion of the Fab’ hybridoma, Fab’DEC205srt was easily isolated from the supernatant with high purity via Ni-NTA gravity flow. To assess whether the Fab’DEC205srt retained antigen specificity, we performed a competitive binding assay. We used a fixed concentration of fluorescently labelled parental NLDC-145 mAb in combination with increasing concentrations of Fab’DEC205srt, unlabeled NLDC-145 mAb, and an isotype control antibody (**Fig. 1F**). Contrary to the isotype control, Fab’DEC205srt and NLDC-145 mAb competed for DEC205 binding with the fluorescent mAb in a dose dependent manner, indicating that the CRISPR engineered Fab’srt binds the same epitope and that antigen specificity is not affected by our CRISPR/HDR approach. The lower EC50 of the Fab’ fragment compared to NLDC-145 is in line with the difference in avidity between monovalent Fab’ and bivalent mAb.

In contrast to conventional proteolytic cleavage to obtain Fab’ fragments, CRISPR/HDR engineering allows the inclusion of various tags. Besides the histag to facilitate purification, we introduced a c-terminal sortag (LPETGG) for chemoenzymatic conjugation using the sortase enzyme (**Fig. S2A**). Unlike classical stochastic chemical conjugation strategies, the sortag facilitates site-specific coupling of cargo to mAbs without the danger of compromising the n-terminal binding region and results in a homogeneous final product (*13, 22*). To show sortag functionality of Fab’DEC205-srt we used a fluorescently labelled sortase substrate probe (GGGCK(FAM), **Fig. S2B**) and (fluorescent) SDS-PAGE analysis. When Fab’DEC205-srt was exposed to 3M sortase (a mutated sortase for higher conversion efficacy) (*23*) alone, the heavy chain c-terminal sortag-histag was hydrolysed resulting in a lower molecular weight heavy chain product (**Fig. 1G, Fig S2C**). However, in the presence of the fluorescent sortase probe the reaction gave quantitative conversion to a fluorescently labelled Fab’DEC205 heavy chain, proving functionality of the sortase recognition motif introduced using our CRISPR/HDR approach.

After successfully converting NLDC-145 to recombinant hybridomas producing fully functional Fab’DEC205-srt, we applied the same strategy to other hybridomas of interest in the field of immuno-oncology. We generated Fab’ hybridomas against murine CD40 (clone FGK45.5), programmed death 1 (PD-1, clone RMP-14) and PD-L1 (clone MIH5). The resulting Fab’ fragments were purified and site-specifically modified to obtain fluorescently labelled Fab’ fragments (**Fig S2C**). Taken together, we demonstrate that our CRISPR/HDR Fab’ hybridoma engineering approach is effective in generating an inexhaustible source of Fab’ fragments equipped with affinity and chemoenzymatic tags with retention of antigen specificity.

### Isotype panel generation via CRISPR/HDR engineering of hybridomas

Encouraged by the success of our one-step CRISPR/HDR strategy to generate Fab’ hybridomas, we continued by further expanding the platform and engineered hybridomas to secrete a diverse set of Fc variants of the same antibody. We designed a CRISPR/HDR genetrap knock-in approach which installs Fc constant domains (heavy chain constant domain 1 (CH1), hinge, CH2 and CH3, sortag, histag, Bsr) upstream of the native rat domains (**Fig. 2A,B**). To force linkage of the genetically engineered foreign CH domains to the native heavy chain variable (VH) domains an artificial splice acceptor was incorporated directly upstream of the new Fc.

**Figure 2.**
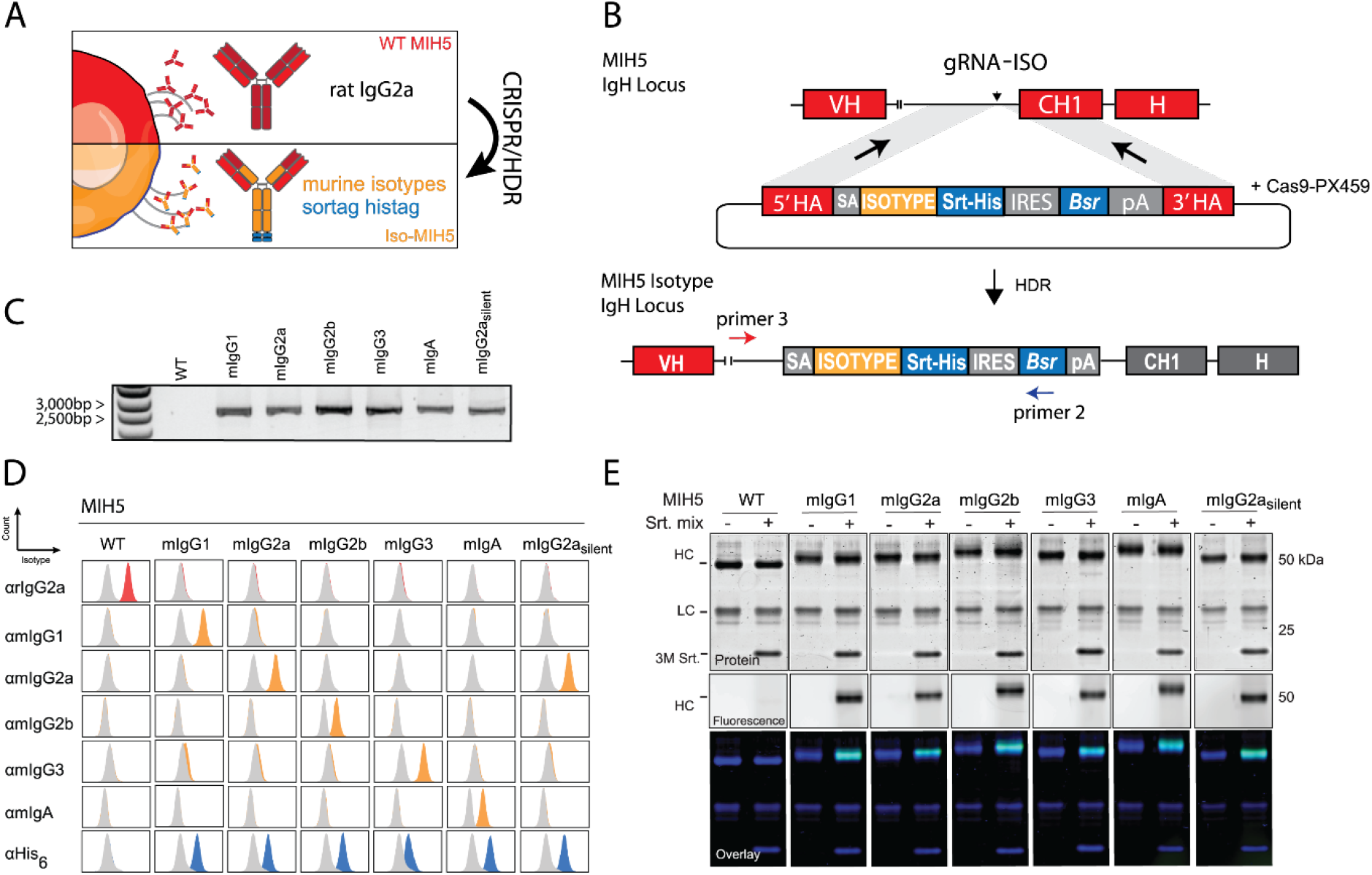
CRISPR/HDR engineering of hybridoma MIH5 to obtain panel of isotype variants. **(A)** CRISPR/HDR strategy to engineer rat IgG2a hybridomas and obtain recombinant hybridomas secreting murine isotypes. **(B)** Shown is the targeted IgH locus of MIH5 with the variable region (VH), constant region 1 (CH1) and hinge annotated. Cas9 is guided by gRNA-ISO to the intronic region upstream of the CH1 to generate a double stranded break. The double stranded break is subsequently resolved via HDR through a donor construct, leading to an in-frame insertion of an splice acceptor (SA, grey), isotype of choice (yellow), a sortag and histag motif (srt-his, blue), an internal ribosomal entry site (IRES), *blasticidin-S deaminase* (*Bsr*), and polyA transcription termination signal (pA) upstream of the native CH1. The insert is enclosed by homology arms (5’HA and 3’HA). **(C)** Three days after electroporation, DNA from CRISPR/HDR targeted MIH5 populations is obtained for PCR with primer 3 and primer 2 (fig 1b). Agarose gel of the PCR product reveals amplicons of the correct size exclusively for the CRISPR/HDR targeted populations. **(D)** After selecting monoclonal hybridoma for each isotype, the supernatant of each isotype modified clone was incubated with PD-L1 expressing target cells (CT26). Displayed plots demonstrate that supernatants exclusively contains the engineered isotype variant with a c-terminal histag, while the original rat IgG2a mAbs is absent. **(E)** Purified MIH5 isotype variants were effectively labeled with the fluorescent probe GGGCK(FAM) using sortase mediated ligation.

To test our method we genetically engineered the hybridoma MIH5, which secretes rIgG2a mAbs targeting mouse immune checkpoint PD-L1, to produce chimeric murine IgG1 (mIgG1), mIgG2a, mIgG2b, mIgG3 and mIgA antibodies. Furthermore, we included a mutant mIgG2a isotype with the amino acid substitutions L234A/L235A/N297A (mIgG2a_silent_), which is described to have reduced affinity for murine FcγRs (*24*). After selection of an optimal sgRNA targeting a PAM sequence 100bp upstream of the rIgG2a CH1 (gRNA-ISO), MIH5 cells were co-transfected with a Cas9 vector containing gRNA-ISO and the panel of isotype HDR donor constructs (**Table S2**). Next, we assessed on-target knock-in integration of blasticidin selected cells by genomic PCR. Analysis showed the presence of PCR products of expected size (∼2600bp, **Fig. 2C**). Monoclonal cell lines were obtained and supernatant was used for flow cytometric screening of the secreted mAbs. CT26 (a mouse colon carcinoma cell line) cells were stimulated with IFN-γ to induce upregulation of PD-L1 before incubation with the supernatants, followed by staining with secondary antibodies for each specific isotype (data not shown). Multiple chimeric clones that expressed the genetically engineered isotypes were identified. We selected a single clone for each isotype and validated the correct knock-in of the isotype of choice in the IgH locus by sequencing. Detailed flow cytometry analysis of the mAbs produced by the selected clones using IFN-γ stimulated CT26 cells revealed that the engineered chimeric mAbs were positive for their respective murine isotypes as well as histag, but not for the native rIgG2a isotype (**Fig. 2D**). To validate antigen specificity of our MIH5 derived engineered antibodies, PD-L1 knock out CT26 cells (CT26^PD-L1^ ^KO^(*25*)) were exposed to the purified mAbs. Contrary to the WT CT26 cells, IFN-γ stimulated CT26^PD-L1^ ^KO^ cells were not stained by our mAbs, proving retention of target specificity (**Fig. S3A**). Western blot analysis of the purified antibodies confirmed substitution of the native rat heavy chain by the genetically introduced mouse heavy chain on protein level (**Fig. S3B**). Notably, if the native splice acceptor adjacent to the CH1 of the rIgG2a is kept in place after CRISPR/HDR, we observed native rIgG2a isotype in the supernatant of chimeric hybridomas as well as in Ni-NTA gravity flow purified proteins, suggesting rat-mouse heavy chain heterodimer mAb formation. We postulated that in these cases, gene elements downstream of the inserted murine heavy chain could still be transcribed due to incomplete transcription termination, facilitating splicing of the VH region to the native rIgG2a isotype. Removal of the native splice acceptor in the HDR donor construct completely abrogated the production of the rat IgG2a isotype (**Fig. S3C**). We equipped all our engineered chimeric mAbs with a c-terminal sortag to facilitate chemoenzymatic functionalization. To confirm functionality of c-terminal sortag we performed a sortase ligation with the fluorescent peptide GGGCK(FAM) (**Fig. 2E**, **Fig. S4**). Indeed, fluorescent SDS-PAGE analysis showed that all the heavy chains from chimeric mAbs were fluorescently tagged, whereas the parental MIH5 antibody was not. Modification of the isotype using our CRISPR/HDR genomic engineering approach is not restricted to the MIH5 hybridoma, as we were able to successfully perform the same strategy on the NLDC-145 hybridoma and obtained a panel of DEC-205 isotype variant mAbs (**Fig. S5**).

We extended the molecular characterization of our engineered mAbs by high-resolution native mass spectrometry analysis of the mIgG2a_silent_. Asparagine (N) on position 297 (N297) is a conserved glycosylation site and is one of the major determinants of FcγR engagement, and hence of therapeutic efficacy of certain mAbs. In mIgG2a_silent_, the N297A amino acid substitution is introduced, rendering it Fc-silent(*24*). To validate the absence of glycans in this engineered mutant, we treated MIH5 WT and mIgG2a_silent_ with the glycosidase PNGase F to remove existing glycans. Subsequently, we performed high resolution native mass spectrometry on treated and untreated samples to assess whether glycans were removed. For the WT samples clear shifts in protein mass were observed, with a leading loss of 2889.6 Da, corresponding to the loss of two glycans (G0F: asialo-agalacto-core-fucosylated biantennary complex-type N-glycan, theoretical mass 2890 Da per mAb, **Fig. 3A**). In contrast, no difference in mass between the PNGase F treated and untreated samples of mIgG2a_silent_ could be measured, indicating the absence of glycans (**Fig. 3B**). Native mass spectrometry analysis of MIH5 mIgG2a variant before and after PNGase F treatment showed a mass difference of 2889.2 Da, corresponding to two G0F glycans, analogous to the WT MIH5 mAb (**Fig. S6**).

**Figure 3.**
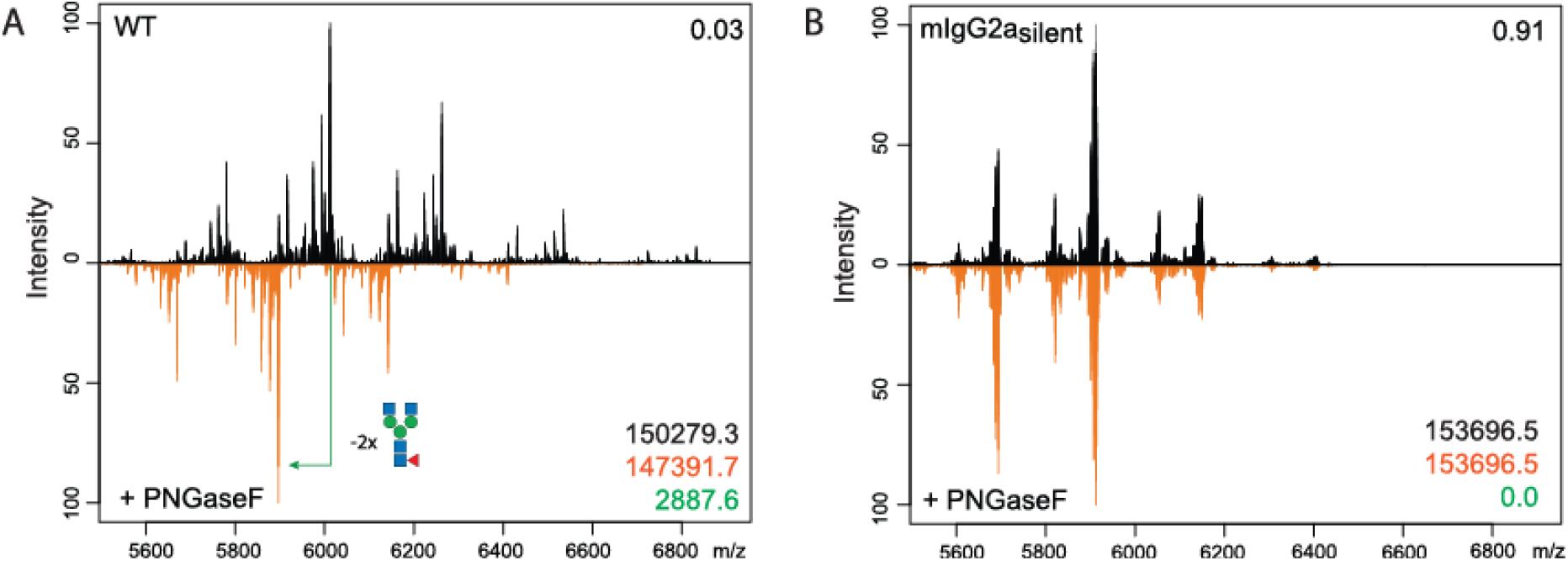
Glycosylation profiling of MIH5 WT and Fc silent variant using high-resolution native mass spectrometry. Purified samples of WT MIH5 **(A)** and mIgG2a_silent_ **(B)** were treated overnight with PNGaseF (orange) to remove existing glycans. Subsequently these samples were compared to untreated samples (black) via high resolution native mass spectrometry. The Pearson correlation coefficient between the two spectra over all ion signals is given in the upper right corner. The molecule mass of the untreated (black) and treated (orange) sample and difference in leading mass (green) is given in the lower right corner.

Our data demonstrates the effective one-step generation of stable hybridomas producing Fc engineered chimeric mAbs using our CRISPR/HDR approach. These mAbs retain their antigen specificity and the addition of affinity and chemoenzymatic tags adds an additional layer of flexibility and utility.

### Biochemical and functional characterization of the engineered antibodies

Having established and validated our hybridoma CRISPR/HDR genomic engineering approach we set out to characterize the functional properties of our isotype-switched mAbs. We first evaluated the binding affinities of our panel of chimeric anti-mouse PD-L1 mAbs for immobilized mFcγRIa, mFcγRIIb and mFcγRIV on a multiplex biosensor via surface plasmon resonance (SPR). We monitored the association and dissociation over time (**Fig. 4A**) and generated affinity plots to calculate the dissociation constants (K_D_) for each receptor concentration. Subsequently, these were interpolated to calculate the affinity of each antibody at receptor concentration giving R_max_=500, as described before (*26*) (**Fig. 4B**). The obtained affinities of mIgG1, mIgG2a and mIgG2b for the FcγRs fall well within the range of previously reported K_D_ values for recombinantly produced isotypes (**Table S3**) (*26*). In summary, mIgG1 did not bind mFcγRI and mFcγRIV, but binds to mFcγRIIb; mIgG2a displayed binding to all tested mFcγRs with highest affinity for mFcγRI, followed by mFcγRIV and mFcγRIIb; mIgG2b binds to mFcγRIIb and mFcγRIV, with stronger affinity for the latter. mIgA and the Fc-silent mutant mIgG2a_silent_ did not show affinity for any of the mFcγRs. The WT rIgG2a displays a FcγR engagement profile similar to the one of mIgG1, binding to the FcγRIIb, but not to FcγRI and FcγRIV. Due to protein aggregation, a characteristic previously reported for mIgG3 (*27, 28*), we were not able to determine the K_D_ values for this isotype variant. Based on the differential affinity for activating and inhibiting receptors reported here, we predicted the capacity of CRISPR engineered mAbs to evoke antibody dependent cellular cytotoxicity (ADCC) (**Fig. 4C**), as others have done in the past (*8*).

**Figure 4.**
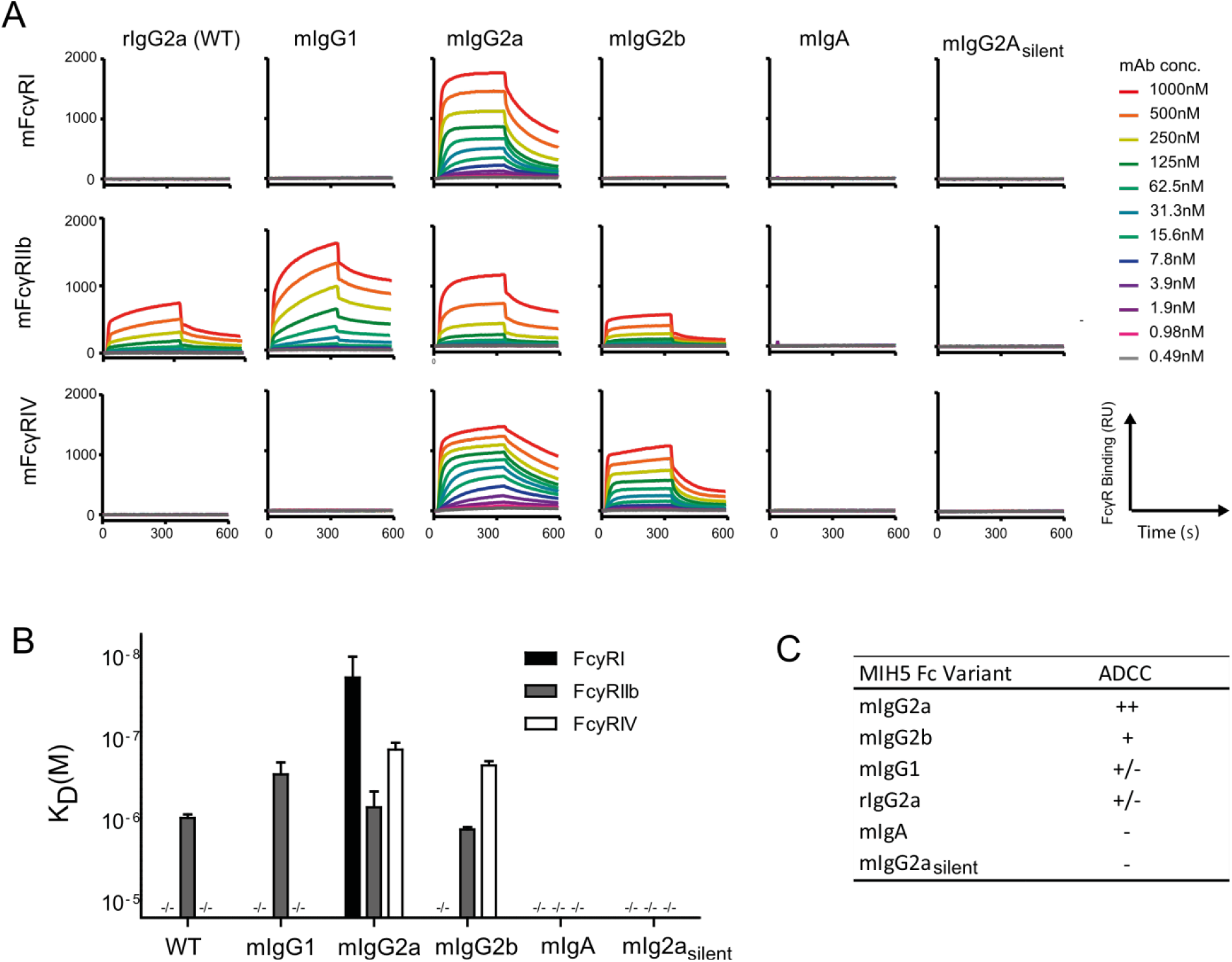
FcγR engagement of MIH5 isotype variants. Representative sensograms **(A)** display interactions of MIH5 WT and MIH5 engineered mAbs (mIgG1, mIgG2a, mIgG2b, mIgA and mIgG2a_silent_) for immobilized mFcγRI, mFcγRIIb and mFcγRIV at increasing concentrations. SPR was performed on four different concentrations of immobilized FcγRs (1nM, 3nM, 10nM and 30nM) to determine affinity (K_D_(M)) of the Fc variants for the different Fc receptors (**B)**, n=3, mean ± s.e.m. **(C)** Predicted ADCC activity for each murine isotype variant on the basis of their differential affinity for FcγRs.

To conclude the characterization of the generated istoype variants we assessed their capacity to induce ADCC in vitro and in vivo. In vitro, we labelled MC38 (murine colon adenocarcinoma) cells with chromium-51, opsonized these with MIH5 Fc variants, and added whole blood from C57BL/6 mice for four hours. To assess specific lysis we measured the chromium-51 release and used an aspecfic mAb (AZN-D1) as a baseline **(Fig. 5A)**. In vivo, we assessed the ability of MIH5 Fc variants to deplete B-cells **(Fig. 5B)**. We labelled isolated splenic B-cells from C57BL/6 mice with Far Red tracer dye and opsonized these with mIgG2a, mIgG2b or mIgA. Simultaneously, we used mIgG2a_silent_ to opsonize splenic B-cells labelled with Violet Blue tracer dye. Subsequently, we mixed the populations in equal ratio and injected these intravenously into LPS-primed C57BL/6 mice. After 24 hours, we sacrificed the mice and isolated the spleens to determine the ratio between the Violet blue and Far red population to assess relative B-cell depletion via flow cytometry (**Fig. 5C**, **Fig S7**). Both in vitro and in vivo assays yielded similar results; mouse IgG2a displayed the strongest capacity to induce Fc mediated killing of target cells, followed by mIgG2b, which is in line with literature and the observed FcyR binding affinities (**Fig. 4A-C**). Neither wildtype rIgG2a, nor its CRISPR engineered variants mIgG1, mIgG3, mIgA and mIgG2a_silent_ were able to mediate specific killing or depletion. As mIgG3, mIgG2a_silent_ and mIgA do not interact with murine Fc receptors, these are unable to mediate ADCC. The lack of effect for rIgG2a and mIgG1 Fc variants can be explained by their ability to bind only a single activating FcyR (FcyRIII), whereas mIgG2a and mIgG2b bind respectively three and two activating FcγRs. Taken together, our in vitro and in vivo data indicate that chimeric and mutant mAbs through hybridoma engineering approach exhibit the same biochemical and immune effector characteristics as their natural and recombinantly produced counterparts.

**Figure 5.**
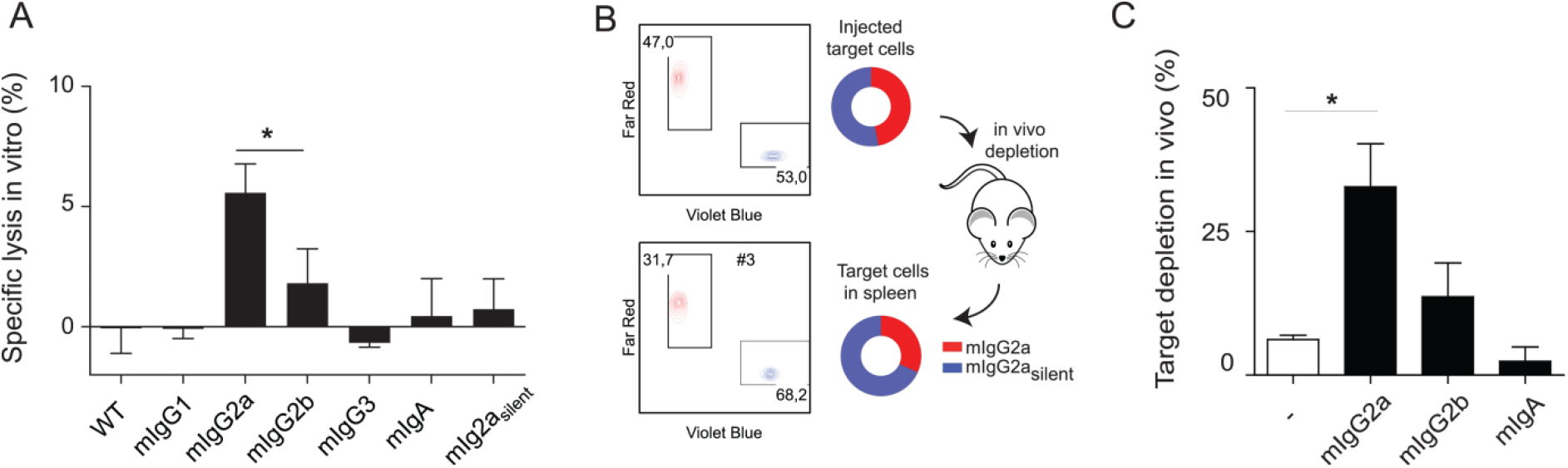
Effector function of MIH5 isotype variants. ADCC in vitro assay **(A)** to compare effector function of MIH5 isotype variants. MC38 cells were ^51^Cr labelled, opsonized with different MIH5 isotype variants, and exposed to whole blood from C57/Bl6 mice for 4 hours. Specific lysis was quantified by measuring ^51^Cr release. *n*=3, mean ± s.e.m (\**p*<0.05). **(B)** Experimental set-up of in vivo depletion assay. Splenic B-cells labelled with violet and red tracer dye were used as target cells for in vivo depletion. The violet B-cells were opsonized with mIgG2a_silent_ variant, while the red B-cells were opsonized with mIgG2a, mIgG2b, or mIgA variant. Subsequently, B-cells were mixed 1:1 (upper plot and diagram) and injected intravenously into C57/Bl6 mice. Twenty-four hours later the spleens were isolated and the ratios between violet and red B-cells determined via flow cytometry (lower plot and diagram). The ratios before injection and after spleen isolation were used to quantify the isotype-specific depletion of the target cells. **(C)** Representative experiment indicating mIgG2a, mIgG2b and mIgA specific depletion of target cells with mIgG2a_silent_ opsonized cells as reference population; *n*=3, mean ± s.e.m (\**p*<0.05).

## Discussion

Here we report the development of a versatile platform for easy and rapid generation of panels of Fc engineered antibodies starting from their parental hybridomas. Using a one-step CRISPR/HDR approach we obtained hybridomas producing either monovalent Fab’ fragments, Fc isotype variants from a foreign species (from rat-anti-mouse to mouse-anti-mouse), and Fc silent mutants without effector functions, within a period of 21 days. Additionally, our method allows incorporation of useful protein tags for purification and site-specific conjugation purposes. Extensive characterization of the produced antibodies revealed that target specificity is uncompromised, and that the CRISPR/HDR engineered chimeric antibodies display their intrinsic Fc-effector functions. Currently, most researchers rely mainly on recombinant production methods to evaluate monoclonal antibodies in different Fc formats. To do so, the variable heavy chain and light chain need to be sequenced, cloned into the appropriate backbones and transfected multiple times into mammalian expression systems to obtain sufficient antibody yields. This process often requires extensive optimization, expertise, knowledge and dedicated resources, not available in most laboratories. In contrast, the CRISPR/HDR strategy outlined here, offers a simple alternative approach requiring a single electroporation step to obtain an unlimited source of target antibody in the isotype format of choice.

Other investigators also described approaches to modulate heavy chain loci in hybridomas via genomic engineering. However, these approaches are either restricted in their engineering capabilities or require multiple sequential genome editing events. For example, Cheong et al. used viral CRISPR-Cas9 vectors to engineer isotype switched mIgG1 and mIgG3 mAbs from IgM-producing hybridomas exploiting the class-switch recombination machinery present in hybridomas (*10*). Although elegant, the technique is restricted to the native heavy chain loci within the target hybridoma. Consequently, generation of a complete isotype panel is restricted to scarcely available IgM-producing hybridomas. Furthermore, placement of recombinant tags, production of mutated isotypes with modulated effector functions or isotypes from a foreign species to generate chimeric mAbs is impossible with this technique. Hashimoto et al. described a multistep method to introduce Fc variants with amino acid substitutions via recombinant mediated cassette exchange (RMCE) (*29*). However, this procedure relies on the prior incorporation of flanking Cre/lox sites required for RMCE, limiting the broad application in other hybridomas. Our single step CRISPR/HDR method is easily adaptable to any hybridoma regardless of background species or isotype to obtain complete panels of engineered antibodies in the format of choice.

We demonstrate, for the first time, the ability to modulate the antibody isotype in hybridomas to a panel ranging from the activating Fc variant mIgG2a, to the Fc-silent mIgG2a_silent_. It is widely recognized that mAb isotype is of major importance for the outcome of antibody-based therapies, especially in the field of cancer immunotherapy. Various studies have shown that the therapeutic efficacy of mAbs directed against immune checkpoints such as CTLA-4, CD25, PD-L1, GITR, and OX-40 depends on their isotype (*4, 5, 30*–*32*). This is mainly due to the interaction of the Fc-region with different components of the immune system, such as FcRs on the surface of immune cells and the C1q protein of the complement system. Substantial efforts have been made to modulate Fc affinity for these different components making use of amino acid substitutions in the Fc-region. This has led to a wide range of described mutants which can be used to increase affinity for activating or inhibitory FcRs (*33*–*35*), ablate Fc-effector functions (*36*), or recruit complement (*37, 38*). Moreover, amino acid substitutions can also be used to control antibody half-life (*39*) or to construct bispecific antibodies (*40, 41*). All of these mutant Fc variants can be implemented in our versatile single step CRISPR/HDR platform.

This method is accessible for any laboratory capable of hybridoma cell line culturing, thus providing a molecular toolbox for repurposing the wealth of established hybridoma cell lines available. Since many hybridoma products are extensively used in preclinical in vivo studies, our method will empower preclinical antibody research by enabling screening of large panels of antibody variants early on in therapeutic antibody development. Moreover, the freedom to add affinity tags and amino acid recognition sequences for chemoenzymatic modification adds an additional layer of flexibility and utility. The ability to site-specifically functionalize the engineered antibody products will find application in the fields of biomolecular engineering, chemical biology, antibody-drug conjugates, multimodal imaging and nanomedicine. We are convinced that this method will open up antibody engineering for the entire scientific community.

## Materials and Methods

### General culture conditions

The following hybridoma clones were genetically engineered for expression of Fab fragments or murine isotypes: NLDC-145 (ATCC®, HB-290™), FGK-45, MIH5, RMP1-14. Other cell lines used in this study were CT26.WT (ATCC® CRL-2638™), CT26.PD-L1^KO^ (*25*), MC38, JAWSII (ATCC® CRL-11904™). For cultivation NLDC-145, FGK-45, MIH5, RMP1-14, CT26, CT26.PD-L1^KO^ and MC38 were grown in complete medium formulation, consisting of RPMI 1640 medium (Thermo Fisher, 11875093), 2mM ultraglutamine (Lonza, BE16-605E/U1), 50 μM Gibco 2-Mercaptoethanol (2-ME) (Thermo Fisher, 21985023), 1x antibiotic-antimycotic (anti-anti) (Thermo Fisher 15240062) and 10% heat inactivated fetal bovine serum (FBS). The cell line JAWSII was cultured according to manufacturer’ protocol. For PD-L1 expression MC38 and CT26 were stimulated overnight with 100ng/mL of recombinant murine IFN-γ.

### Cloning of CRISRP-Cas9 and donor constructs

The genomic sequence of the rat IgG2a heavy chain locus was identified via the Ensembl rate genome build Rnor_6.0 (accession: AC_0000741) and used for the design of the HDR donor templates. gRNA-H (CTTACCTGTACATCCACA) and gRNA-ISO (CAAGTCTGTGGCTGTCTACA) were designed using the CRISPR tool from the Zhang lab (http://crispr.mit.edu) and ordered as single stranded oligos with from Integrated DNA Technologies (IDT) with the appropriate overhangs for cloning purposes. The oligos were phosphorylated with T4 PNK enzyme by incubation at 37°C for 30 minutes and annealed by incubation at 95°C for 5 minutes followed by gradually cooling to 25°C using a thermocycler. The annealed oligos were cloned into the plasmid pSpCas9(BB)-2A-Puro (PX459), obtained as a gift from Feng Zhang (Addgene plasmid #62988). All CRISPR/Cas9 and HDR constructs were purified with nucleobond® Extra midi kit (Machery-Nagel, 740410.100) according to manufacturer’ protocol. Synthetic gene fragments containing homologous arms and desired inserts were obtained via IDT and cloned into the PCR2.1 TOPO TA vector (ThermoFisher, K-450001). Removal of the native splice acceptor in the 3’ homologous arm of HDR templates was achieved via site-directed mutagenesis (Agilent Technologies, 200523).

### Hybridoma nucleofection with donor constructs and CRISPR-Cas9

Nucleofection of the HDR template and CRISPR-Cas9 vectors was performed with SF Cell Line 4D-Nucleofection X Kit L (Lonza, V4XC-2024). Before nucleofection hybridoma cells were assessed for viability and centrifuged (90*g*, 5 minutes), resuspended in PBS/1% FBS and centrifuged again (90*g*, 5 minutes). One million cells were resuspended in SF medium with 1 μg of HDR template and 1 μg of CRISPR-Cas9 vectors or 2 μg of GFP vector (control) and transferred to cuvettes for nucleofection with the 4D-Nucleofection System from Lonza (Program SF, CQ-104). Transfected cells were transferred to a 6-well plate in 4 mL of complete medium. The following day the cells were transferred to a 10 cm petridish in 10 mL of complete medium, supplemented with 10-20 μg/mL of blasticidin (Invivogen, anti-bl-05). Antibiotic pressure was sustained until GFP transfected hybridomas were dead and HDR transfections were confluent (typically between day 10-14). Cells were subsequently clonally expanded by seeding the hybridomas in 0,3 cells/well in u-bottom 96-well plates in 100 μL of complete medium. After one week, supernatant from wells with a high cell density were obtained for further characterization.

### Genomic DNA isolation, PCR and sequencing

One week after nucleofection with HDR and targeting constructs a minimum of 10,000 cells of the surviving cell population were collected for genomic phenotyping. DNA was extracted using the genomic DNA isolate kit from Bioline (BIO52067). The DNA pellet was resuspended in ultra-purified water and used for PCR analysis. For PCR confirmation of fab’ hybridoma we used primer 1 (TGTAGGAGCTTGGGTCCAGA) which hybridizes upstream of the 5’ homolog arm of the HDR template and primer 2 (ATACATTGACACCAGTGAAGATGC) which hybridized with the BSD gene. For confirmation of the integration of isotype constructs we used primer 3 (GGCGACCTGTAACAACTTGG), annealing upstream of the 5’ end of the HDR and primer 2. PCR products were visualized on a 1% (wt/vol) agarose gel containing Nancy-520 dye stain. DNA was excised, purified (Macherey-Nagel, 740609) and validated via Sanger sequencing.

### Flow cytometry

Typically ten days after limiting dilution, supernatants from wells with high cell density were transferred to wells containing 20,000 target cells and incubated for 20 minutes at 4°C in a V-bottom 96-well plate. Subsequently, plates were centrifuged (90*g*, 2 minutes) and hybridoma supernatant discarded by flicking. Plates were washed twice with PBS/5% FBS and incubated with anti-rat IgG2a (PE) (Thermofisher, 12-4817-82) or anti-histag (PE) (Biolegend, 362603) to assess fab’-secreting hybridomas. For isotype grafted hybridomas, the isotype was determined by using the following panel of secondary antibodies on histag positive supernatants: anti-mouse IgG1 (PE) (Thermofisher, 12-4015-82), anti-mouse IgG2a (PE) (Thermofisher, 12-4817-82), anti-mouse IgG2b (FITC) (11-4220-82), anti-mouse IgG3 (A488) (Thermofisher, A21151), anti-mouse IgA (FITC) (11-4817-82). To assess specificity 50,000 JAWSII cells/well were seeded in a V-bottom 96 well plate and stained with Fab’ DEC2 05, NLDC-145 or AZN-D1 (αDC-SIGN) in concentrations ranging between 0.01 – 1000nM. After 10 minutes of incubation at 4°C, 6.7nM of NLDC-145 (PE) (Biolegend, 138213) was added to the wells, after 30 minutes plates were washed and acquired on FACS.

### Western Blot

Supernatants from fab’ engineered hybridomas were analyzed for presence of fab’ fragments by anti-histag staining and anti-rat IgG (H+L) staining. The purified murine MIH5 isotypes were stained for presence of rat and histagged heavy chains. Samples were run under reducing conditions on 12% sodium dodecul sulfate polyacrylamide gel electrophoresis (SDS-PAGE). Gels were transferred on PVDF membrane and blocked with 3% BSA in PBS-Tween 0,02% (PBS-T) and stained with rabbit anti-histag antibody (abcam, ab137839), goat anti-rabbit IgG (H+L) (IRD800) (LI-COR, 926-32211) and goat anti-rat IgG (H+L) (AF680) (Thermofisher, A-21096).

### Antibody production and isolation

Engineered hybridomas selected for production were expanded in T175 flasks containing 20mL of complete medium. Once confluent, hybridomas were cultivated for another 1-10 days in RPMI-1640 supplemented with 2mM ultraglutamine, 1x AA, 50uM 2-ME, and 10% FBS. For larger antibody titers, 20 million cells were seeded in Celline Bioreactor Flasks (Argos, 900-05) and cultivated for 1 week. Antibody-containing-supernatants were separated from cells via centrifugation (90*g*, 5 minutes), filtered through a 20μM filter and supplemented with 10mM imidazol (Sigma-Aldrig, I2399). Subsequently the hybridoma supernatant was run over a Ni-NTA column (Qiagen, 30210). The column was subsequently washed with 10 column volumes of PBS with 25mM imidazole before antibody elution with PBS containing 250mM imidazole. Buffer exchange to PBS (for mIgG3; 300mM NaH_2_PO_4_, 50mM NaCl_2,_ pH 6.5) was performed via ultracentrifugation with amicon Ultra-15 Centrifugal filter units (Sigma, Z717185). Antibodies from WT hybridomas were obtained by cultivating parental hybridoma using CD Hybridoma medium (Thermofisher, 11279023) supplemented with 2mM ultraglutamine, 1x AA and 50uM 2-ME. Antibodies were purified from medium using protein G gravitrap (Sigma, 28-9852-55) according to manufacturer protocol. Antibody concentration was determined via absorption at 280nm with an extinction coefficient of 1.4. Protein purity was assessed by SDS-PAGE using SYPRO™ Ruby Protein Gel Stain (ThermoFisher, S12000).

### Sortase-mediated ligation

To assess sortase-mediated conjugation a small nucleophilic peptide (GGGCK(FAM)) was constructed via solid-phase-peptide synthesis. Antibodies were dialysed into sortase buffer (150mM Tris, 50mM NaCl, 10mM CaCl_2_, pH7.5) via ultracentrifugation. Sortase mediated reactions were carried out with 5 μg of substrate (mAb or Fab’) in 10 μL volumes, containing 1 (Fab’) or 2 (mAb) molar equivalents of tri-mutant sortase A Δ59 (3M srt) (*23*), and 50 (Fab’) or 100 (mAb) molar equivalents of GGGCK(FAM) peptide. After one hour of incubation at 37°C, the reaction was stopped via addition of EDTA (10mM final concentration). For each reaction, 500ng was loaded on reducing SDS-PAGE and analyzed for fluorescence and protein using SYPRO™ Ruby Protein Gel Stain on Typhoon Trio+.

### Native Mass spectrometry analysis

mAbs were treated overnight with four units of PNGase F (Roche, IN, USA) at room temperature. Subsequently, treated and untreated samples were buffer exchanged into 150 mM aqueous ammonium acetate (pH7.5) by ultrafiltration (vivaspin 500, Sartorius Stedim Biotech Germany) with 30kDa cut-off filter. Sample concentrations were adjusted to 5µM, before 4µL was used for native MS analysis. Native MS analysis was carried out with a modified Exactive Plus Orbitrap instrument with extend mass range (EMR) as described before(*42*). Intact Mass software was used for zero-charge deconvolution of the spectra to determine the antibody mass(*43*).

### FcγR engagement

Surface plasmon resonance measurement were performed on an IBIS MX96 (IBIS technologies) with a similar methodology as described before(*26*). In short, biotinylated mouse FcγRI, FcγRIIb and FcγRIV were immobilized in duplicate, ranging from 30nM to 1nM onto a single SensEye G-streptavidin sensor. Then MIH5 Fc variants were injected over the IBIS in PBS supplemented with 0.075% Tween-80 in concentrations ranging from 0.49nM to 1000nM. K_D_ calculation was performed by fitting a 1:1 langmuir binding model to the RU_360sec_ values at each antibody concentration. For consistency, these fits were performed at each receptor concentration and the final K_D_ values were calculated by interpolating to R_max_=500. The measurements were performed in triplicate.

### ADCC in vitro

Mouse whole blood ADCC with a ^51^Cr release assay was performed as described before(*44*). Briefly, M38 cells were stimulated overnight with 100ng/uL of IFNy to upregulate PD-L1 expression and used as target cells. Subsequently, the target cells (5000 cells/well) were labelled with 100 μCu 51Cr for 2 h at 37°C and washed in complete medium. CRISR/HDR engineered antibodies were added in a final concentration of 10ug/mL, followed by addition of whole blood (25 μL) derived from pegylated G-CSF stimulated C57/Bl6 mice expressing human CD89. Cellular mix was incubated for 4 hours in 200 μL RPMI1640 + 10% FCS at 37°C. ^51^Cr release was counted in 50 ul supernatant with a liquid scintillation counter. Percentage of specific lysis was calculated as follows: (experimental cpm - basal cpm)/(maximal cpm - basal cpm) × 100, with maximal lysis determined in the presence of 5% triton and basal lysis in the present of an antibody control (AZN-D1, αDC-SIGN).

### B-cell depletion in vivo

B-cells were isolated from two C57BL/6 mice via MACS depletion (CD43 microbeads, 130-049-801 MACS) and labelled with Cell Trace Violet (ThermoFisher, C34557) or Cell Trace Far Red (ThermoFisher, C34564). To increase PD-L1 expression, labelled B cells were treated with 100ng/mL IFN-γ and 10ng/mL of IL-4 overnight at 37°C. Following day, 9 million Far red labelled B-cells were incubated with 25µg/mL of MIH5 mIgG2a, mIgG2b or mIgA and 27 million violet labelled B-cells with MIH5 mIgG2a_silent_. Far red labelled B-cells and violet labelled B-cells were mixed 1:1 and intravenously injected into C57BL/6 mice (6 million/mouse), which were stimulated 6 hours earlier with LPS (1µg/g). Part of the cell suspension was analyzed to determine the ratio between Far Red and Blue B-cells injection. Twenty-four hours after injection, isolated spleens were meshed through a 100µm cell strainer using a syringe plunger. The resulting cell suspension was centrifuged (90*g*, 5 minutes) and incubated in ammonium chloride potassium solution to lyse erythrocytes. After 5 minutes of incubation cells were washed with PBS two times and analyzed with flow cytometry to determine the ratio between violet B-cells and far red B-cells. To determine the relative depletion the following calculation was used: 1-(Injected _(Violet:Red)_/Isolated _(Violet:Red)_)x100. Mice were maintained under specific pathogen-free conditions at the Central Animal Laboratory (Nijmegen, the Netherlands) and treated according to guidelines for animal care of the Nijmegen Animal Experiments Committee.

## Supplementary Materials

Figure S1. Genomic map and annotated basepair sequence of rat IgG2a constant domains.

Figure S2. Sortagging of CRISPR/HDR engineered hybridomas derived Fab’ fragments.

Figure S3. Characterization of MIH5 Fc variants hybridomas.

Figure S4. Murine isotype panel generation of NLDC-145 via CRISPR/HDR.

Figure S5. Raw images of sortagging of MIH5 isotype variants

Figure S6. Glycosylation profile of MIH5 mIgG2a via native mass spectrometry

Figure S7. Gating and FACS plots of isotype dependent depletion in vivo Table S1. Fab’ donor construct for HDR.

Table S2. Isotype donor constructs for HDR.

Table S3. FcγR affinity values of MIH5 Fc variants and comparison to literature.

## Acknowledgments

We thank Sander van Duijnhoven (Radboudumc, Nijmegen, the Netherlands) for kindly providing the 3M Sortase A and K.A. Marijt for kindly providing the CT26.PD-L1^KO^ cell line. This work was supported by the Netherlands Organisation for Scientific Research (NWO; Project number 13770) and in part through the proteomics facility Proteins@Work (Project 184.032.201) embedded in the Netherlands Proteomics Centre. C.F. is recipient of Dutch Cancer Society KWO grant 2009-4402, the NWO Spinoza award and ERC Adv. Grant PATHFINDER (269019). M.V. is recipient of ERC Starting grant CHEMCHECK (679921) and a Gravity Program Institute for Chemical Immunology tenure track grant by NWO. T.C. is a PhD student fellow of the Gravity Program Institute for Chemical Immunology. F.A.S. is recipient of an LUMC fellowship.

## Author contributions

M.V. and F.A.S. conceived the project and contributed equally. J.L., A.J.R.H., G.V., C.G.F., M.V. and F.A.S. provided guidance and support. J.M.S.S., J.L., A.H., G.V., M.V. and F.A.S. designed experiments. J.M.S.S., F.L.F., M.V., Y.D., A.C., T.C., J.A.C.B., M.E., A.E.H.B., M.N. and F.A.S. performed the experiments. I.M.H., M.F.F., A.M.D.B., C.M.L.G. and D.D. contributed experimentally. J.M.S.S., M.V. and F.A.S. wrote the paper.

## Competing interests

The authors declare no competing interests.

## Data and materials availability

All material and data used in this study are available from the corresponding authors upon reasonable request.

