## Supplemental information for "Functional diversification of hybridoma produced antibodies by CRISPR/HDR genomic engineering"

**A**

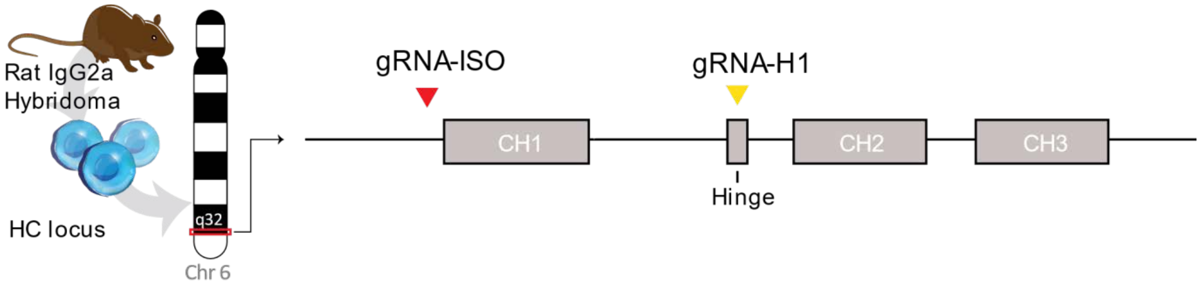


**B**

```
Rattus norvegicus: Rn Celera
Chr.6 (AC_0000741)
129,955,150 – 129,957,604

AGAAAGATCTGAGTAGAACCAAGGTAAAAAGTGTGGGTAAAAACACATGTTCACAGGCCTGGCTGACATGATGCTGGGCACGTATGGAGGCAAA
GTCAAGAGGGCAGTGTAAGGGCCAGAAGTGAATCCTGACCCAAGAATAGAGAGTGCTAAACCTACGTAGATCGAAGCCAACTAAAAAGACAAGC
TACAAAACGAAGCTAAGGCCAGAGATCTTGGACTGTGAAGAGTTCAGAGAACCTAGGATCAGGAACCATTAGTAACAGGCCAAGGAAGATAGAA
GCTGCCTAGGACTTGGCAAGAGCCAACATGGTTGGACTGGAAAAGAAAGGAGGAGACAGAAGACAGGAGAGATGTGCCAACTTGATTTTGGGCT
TCACTGTTGTCCATACTGTGTGCAGCCATATGGCCCACAGATAACAGGTTTAGCCGAGGAACACAGATACCCACATTGGACAATGGTGGGGGAA
CACAGATACCCATACTACAGGGCTCTTTAGGGCATTTCCTGAAAGTGTACTAGGAGTGGGACTGGGCTCAAAGGGATTAGGTGTGATCTGGCCT
GGTGAGGCTGACATTGGCAAGCCCAATGGTTGGGTGTTGCCTCCTCCATGTAGACAGCCACAGACTTGGGGAGGGTACAAAATGGAGGACTTGT
AGGAGCTTGGGTCCAGACCTGTCAGACAAAATGATCACGCATACTTATTCTTGTAGCTGAAACAACAGCCCCATCTGTCTATCCACTGGCTCCT
GGAACTGCTCTCAAAAGTAACTCCATGGTGACCCTGGGATGCCTGGTCAAGGGCTATTTCCCTGAGCCAGTCACCGTGACCTGGAACTCTGGAG
CCCTGTCCAGCGGTGTGCACACCTTCCCAGCTGTCCTGCAGTCTGGACTCTACACTCTCACCAGCTCAGTGACTGTACCCTCCAGCACCTGGTC
CAGCCAGGCCGTCACCTGCAACGTAGCCCACCCGGCCAGCAGCACCAAGGTGGACAAGAAAATTGGTGAGAGAACAACCAGGGGATGAGGGGCT
CACTAGAGGTGAGGATAAGGCATTAGATTGCCTACACCAACCAGGGTGGGCAGACATCACCAGGGAGGGGGCCTCAGCCCAGGAGACCAAAAAT
TCTCCTTTGTCTCCCTTCTGGAGATTTCTATGTCCTTTACACCCATTTATTAATATTCTGGGTAAGATGCCCTTGCATCATGACATACAGAGGC
AGACTAGAGTATCAACCTGCAAAAGGTCATACCCAGGAAGAGCCTGCCATGATCCCACACCAGAACCAACCTGGGGCCTTCTCACCTATAGACC
ATACTAACACACAGCCTTCTCTCTGCAGTGCCAAGGGAATGCAATCCTTGTGGATGTACAGGTAAGTCACTAGGACTATTACTCCAGCCCCAGA
TTCAAAAAATATCCTCAGAGGCCCATGTTAGAGGATGACACAGCTATTGACCTATTTCTACCTTTCTTCTTCATCTACAGGCTCAGAAGTATCA
TCTGTCTTCATCTTCCCCCCAAAGACCAAAGATGTGCTCACCATCACTCTGACTCCTAAGGTCACGTGTGTTGTGGTAGACATTAGCCAGAATG
ATCCCGAGGTCCGGTTCAGCTGGTTTATAGATGACGTGGAAGTCCACACAGCTCAGACTCATGCCCCGGAGAAGCAGTCCAACAGCACTTTACG
CTCAGTCAGTGAACTCCCCATCGTGCACCGGGACTGGCTCAATGGCAAGACGTTCAAATGCAAAGTCAACAGTGGAGCATTCCCTGCCCCCATC
GAGAAAAGCATCTCCAAACCCGAAGGTGGGAGCAGCAGGGTGTGTGGTGTAGAAGCTGCAGTAGGCCATAGACAGAGCTTGACTTAACTAGACT
TCTGCCCTCTTACTGACCTCCATGCTGACCACTCTCTGTATCCACAGGCACACCACGAGGTCCACAGGTATACACCATGGCGCCTCCCAAGGAA
GAGATGACCCAGAGTCAAGTCAGTATCACCTGCATGGTAAAAGGCTTCTATCCCCCAGACATTTATACGGAGTGGAAGATGAACGGGCAGCCAC
AGGAAAACTACAAGAACACTCCACCTACGATGGACACAGATGGGAGTTACTTCCTCTACAGCAAGCTCAATGTAAAGAAAGAAACATGGCAGCA
GGGAAACACTTTCACGTGTTCTGTGCTGCATGAGGGCCTGCACAACCACCATACTGAGAAGAGTCTCTCCCACTCTCCTGGTAAATGATCCCAG
AGTCCAGTGGCCCCTCTTGGCCTAAAGGATGCCAACACCTACCTCTACCACCTTTCTCTGTGTAAATAAAGCACCCAGCTCTGCCTTGGGACCC
TGCAAAAATGTCCTGGTTCTTTCTGAGATACAGAGTCCAGTGAGGTCATGGGCTGAGGGGCATCCAGGGTTTGAGGCCTGAGGTTTGACTAAGG
AAAAAGGGTGG
```

**Figure S1. Genomic map and annotated basepair sequence of rat IgG2a constant domains.** The genomic location **(A)** and annotated basepair sequence **(B)** and of the IgH locus of rat IgG2a located on chromosome 6 (Rn celera, AC_0000741) are given. The exons of CH1, Hinge, CH2 and CH3 are indicated (grey highlight) with splice acceptor and donor sites (underlined, cursive). The targeted protospace adjacent motifs (PAMs) for gRNA-H (yellow) and gRNA-ISO (red) are indicated.

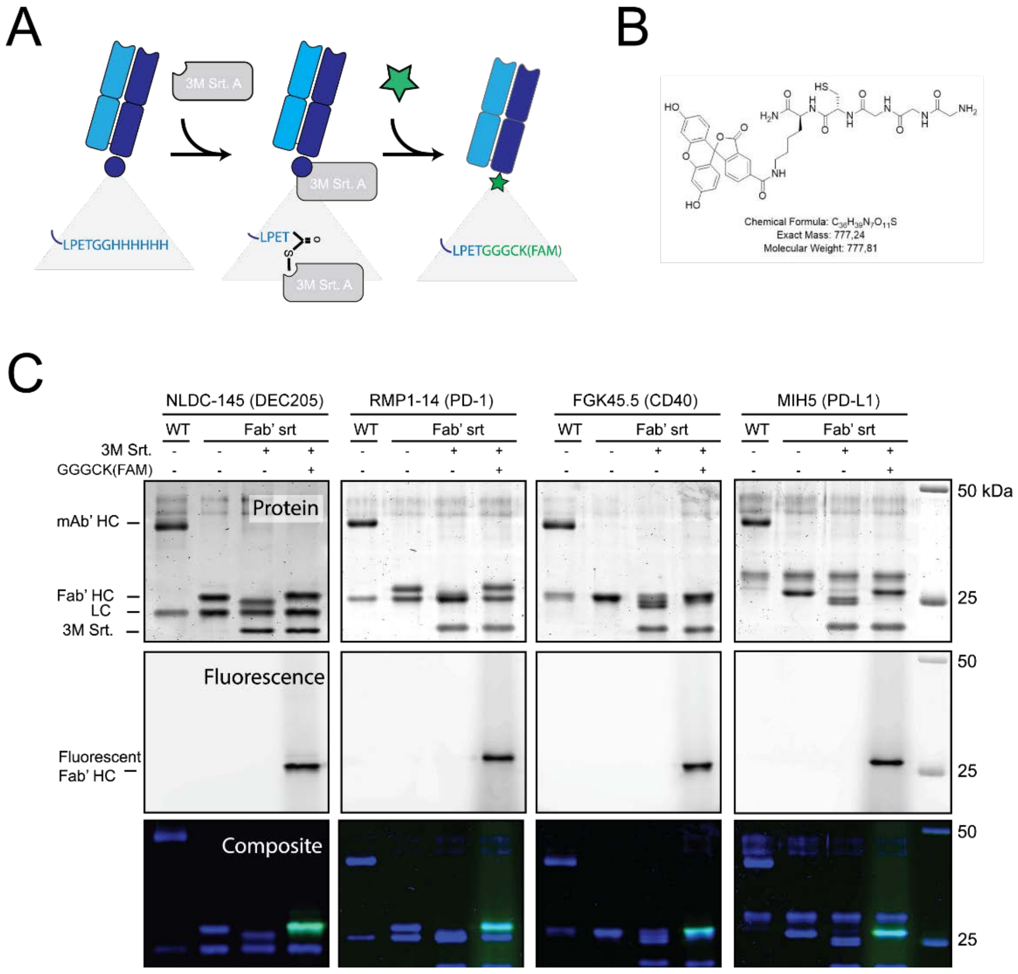


**Figure S2. Sortagging of Fab' fragments derived from CRISPR/HDR engineered hybridomas.** The CRISPR/HDR strategy as outlined in Fig. 1 was also performed on hybridomas RMP1-14 (PD-1), FGK45.5 (CD40) and MIH5 (PD-L1). The resulting Fab' hybridomas secreted all Fab'fragments which could be easily isolated from the supernatant, demonstrating universal applicability of this approach. **(A)** All Fab' fragments were equipped with a c-terminal sortag motif to perform sortase mediated ligation. **(B)** We synthesized the fluorophore GGGCK(FAM) to functionaliz the c-terminus of the Fab' fragment heavy chain. **(C)** To this purpose, we incubated 5µg of Fab' fragment (10µM) with equimolar amounts of an evolved sortase (3M Srt., 10µM) and with or without 50 molar equivalents of GGGCK(FAM) for one hour at 37ºC. From each reaction mix we loaded 5ug on reducing SDS-page together with the parental mAb and unmodified Fab and acquired the protein and fluorescent scans. The left panel displays the protein and fluorescent scans of Fab'DEC205-srt, of which the composite is shown in **Fig. 1G**.

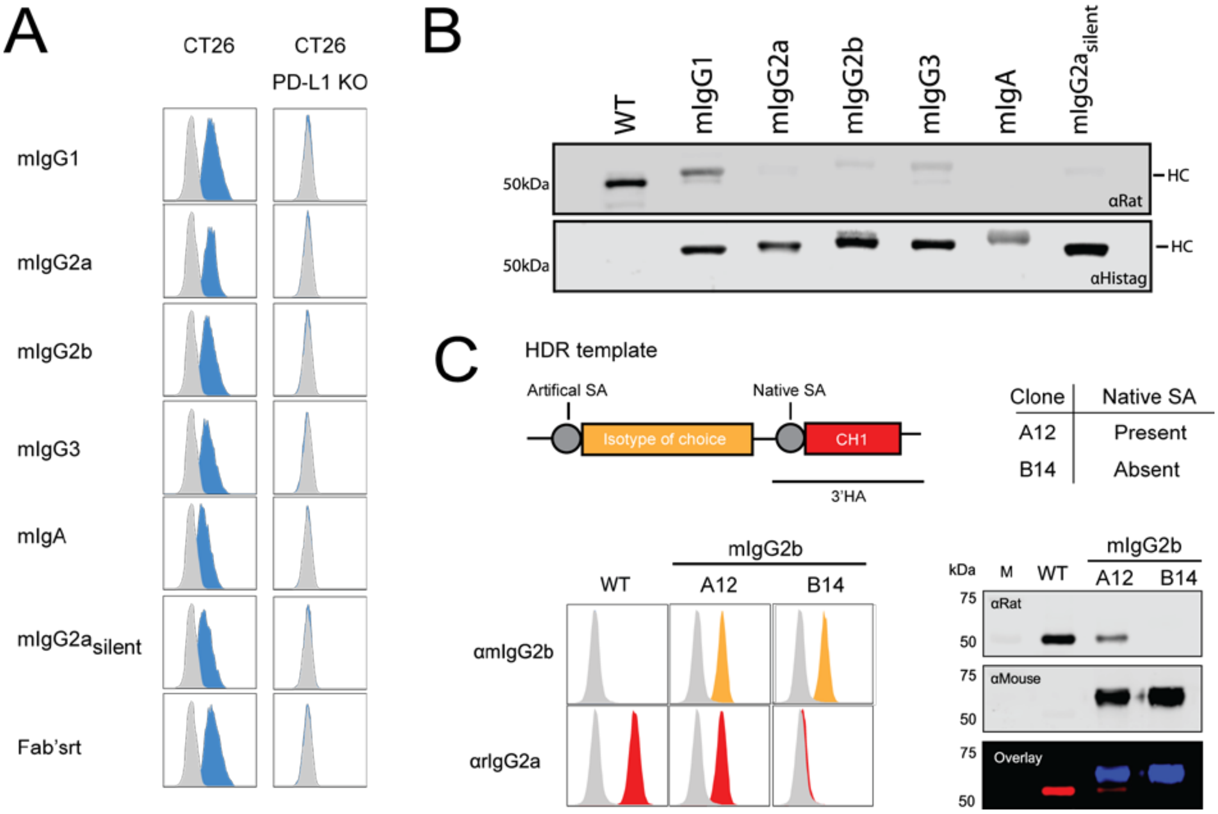


**Figure S3. Characterization of MIH5 Fc variants hybridomas. (A)** FACS plots of CT26 and CT26$^{PDL1\ KO}$ stained with MIH5 Fc variants in combination with αhistag indicate specificity for PD-L1 is retained. **(B)** Western blot analysis demonstrates substitution of the rat HC for murine HC for each MIH5 Fc variant. **(C)** Presence of the original splice acceptor adjacent to the rat CH1 exon results in low secretion of the native rat IgG2a MIH5 in the supernatant (mIgG2b-A12). In the supernatant of engineered hybridomas where the splice acceptor is removed (mIgG2b-B14), production of the native isotype is completely abrogated as determined by flow cytometry and western blot.

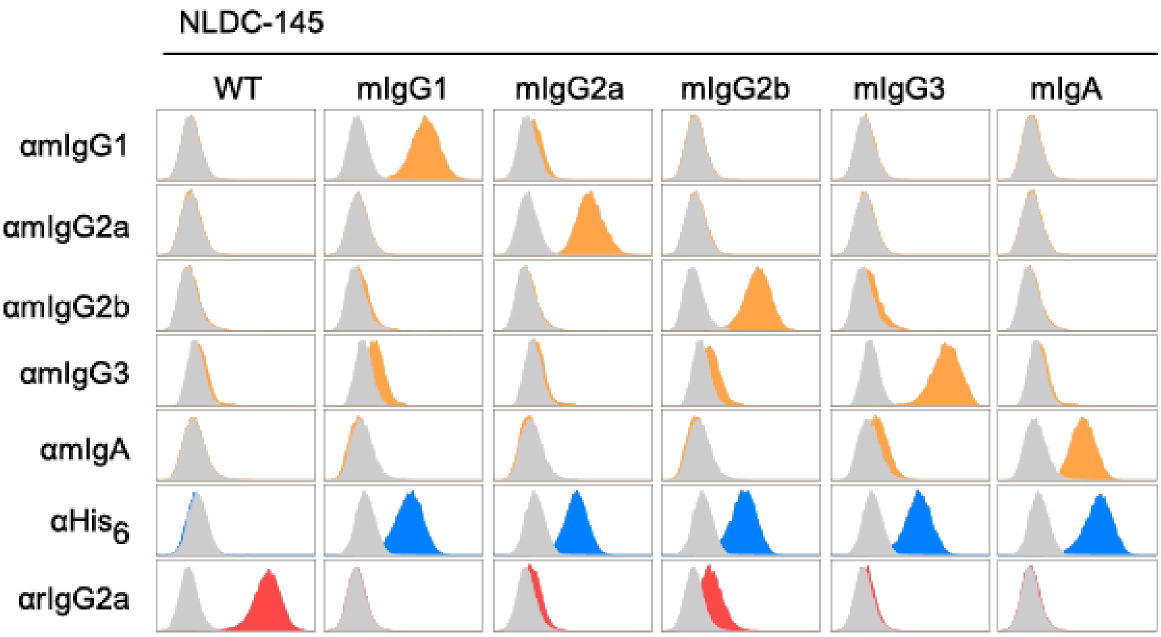


**Figure S4. Murine isotype panel generation of NLDC-145 via CRISPR/HDR.** The CRISPR/HDR strategy as outlined in **Fig. 2** was applied to the NLDC-145 to obtain recombinant hybridomas secreting murine IgG1, IgG2a, IgG2b, IgG3 and IgA. For each isotype, hybridoma supernatant was used to stain DEC-205 expressing cell line JAWS II in combination with a panel of secondary antibodies against rat IgG2a, histag and murine isotypes.

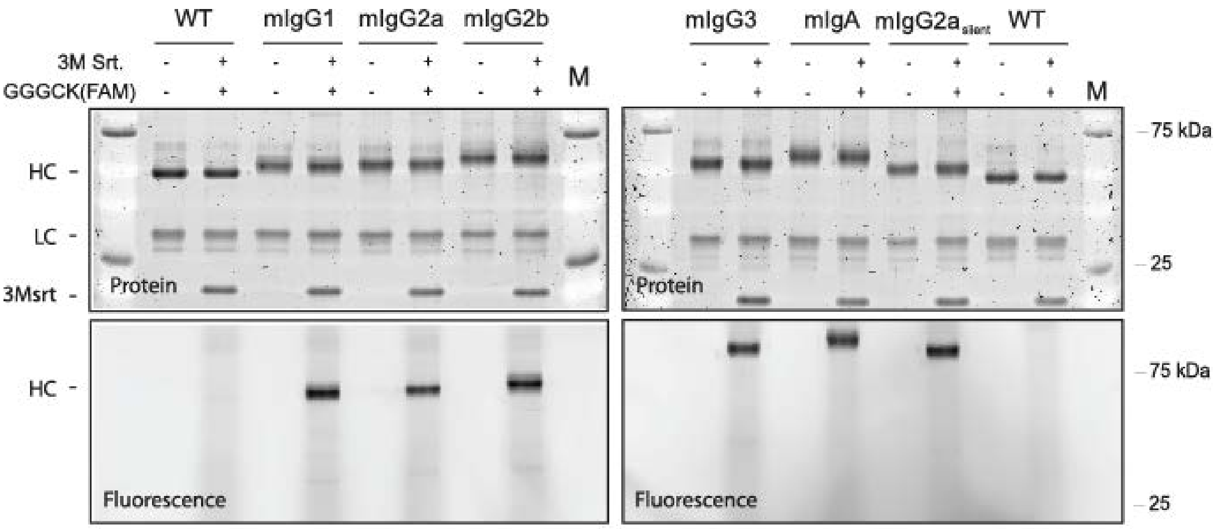


**Figure S5. Raw images of sortagging of MIH5 isotype variants.** Raw images of protein and fluorescent SDS-page scans of sortagging of MIH5 isotype variants as displayed in **Fig. 2E**. Of each antibody 5μg was incubated with 1 molar equivalent of 3m Sortase (3m Srt) and 100 molar equivalent GGGCK(FAM) for one hour. From each reaction 500ng was run on reducing SDS-PAGE. Subsequently, protein and fluorescent images of the SDS-page were acquired.

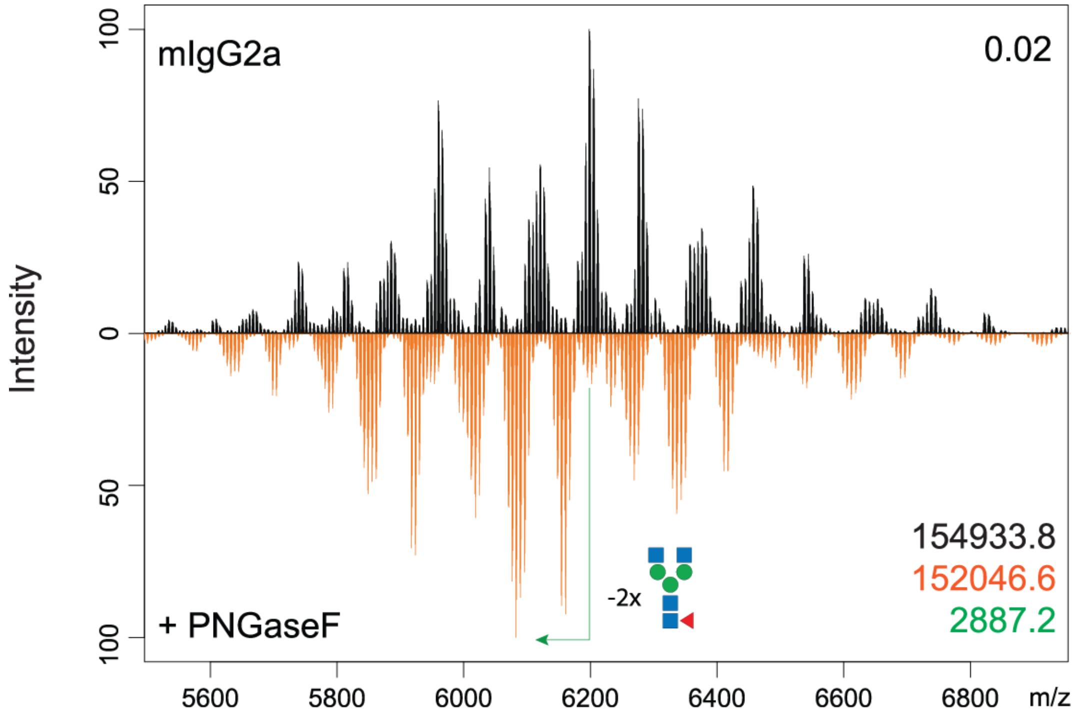


**Figure S6. Glycosylation profile of MIH5 mIgG2a via native mass spectrometry.** Purified MIH5 mIgG2a was treated overnight with PNGAseF (orange) and compared to an untreated sample (black) via high resolution native mass spectrometry. The Pearson correlation coefficient between the two spectra over all ion signals is given in the upper right corner. The molecular mass belonging to the base peak of the untreated (black font) and treated (orange font) sample and the leading difference in mass (green) is depicted in the lower right corner. The difference in mass between treated and untreated samples indicates the loss of two glycans.

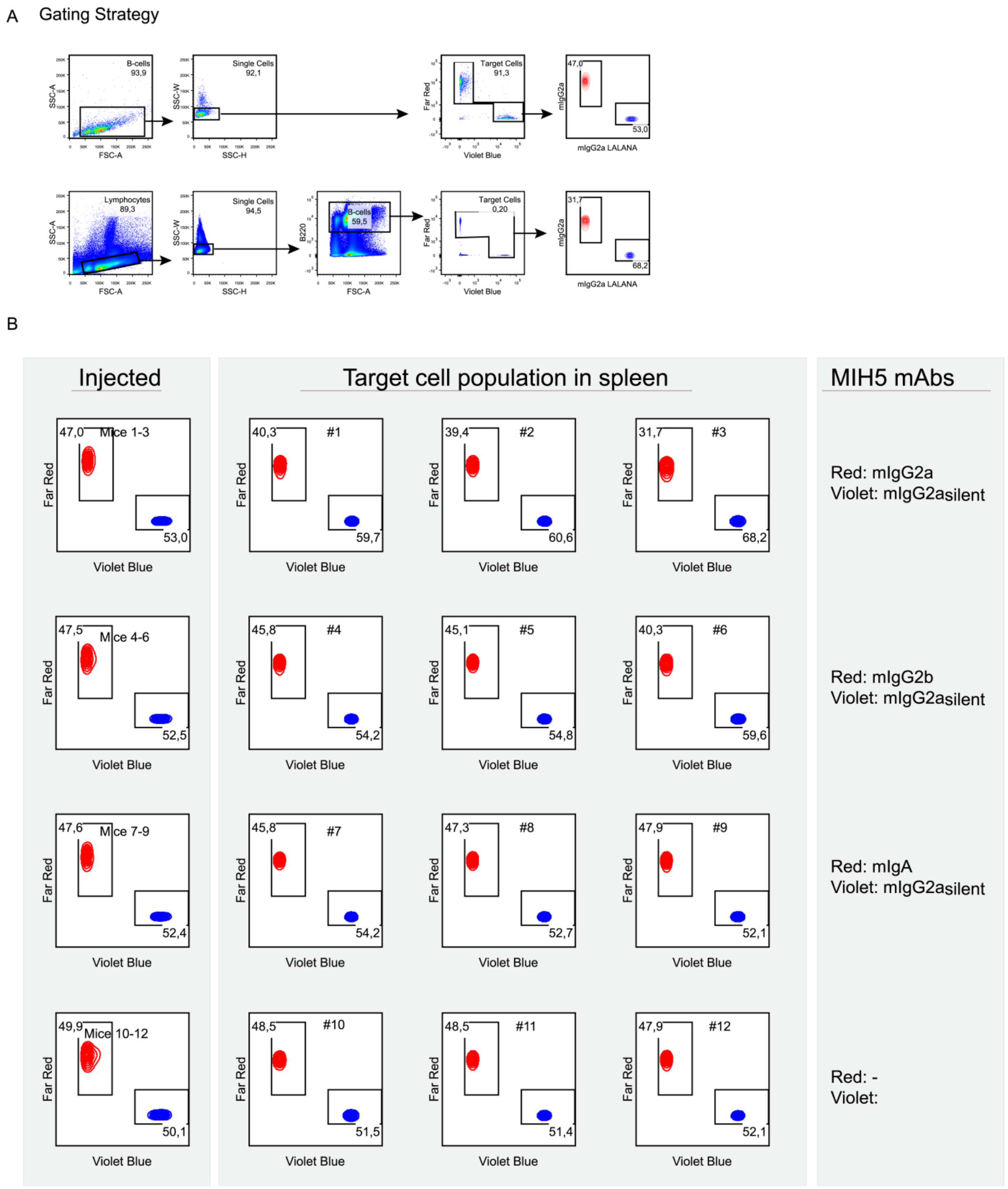


**Figure S7. Gating and FACS plots of isotype dependent depletion in vivo.** Isolated B-cells were stimulated overnight with IFN-y and labeled with either red or violet tracer dyes. Subsequently, the violet B-cells were opsonized with MIH5 mIgG2a$_{silent}$, while the red B-cells were opsonized with MIH5 mIgG2a, mIgG2b or mIgA. The populations were mixed and intravenously injected into LPS stimulated BL/6 mice. After 24 hours the mice were sacrificed and spleens isolated to determine the ratio of violet blue and far red target cells. These were used to determine the isotype dependent target cell depletion. The gating strategy (**A**) and the plots (**B**) depicting the labeled B-cell ratios used for determing the specific target depletion in **Fig 5C** are given.

**Table S1. Fab' donor construct for HDR.** Table displays sequences of each feature of the donor construct used to convert rat IgG2a hybridomas to sortagable Fab'fragment secreting cell lines.

| Isotype | Sequence |
|---|---|
| 5' HA | CCTGGAACTCTGGAGCCCTGTCCAGCGGTGTGCACACCTTCCCAGCTGTCCTGCAGTCTGGACTCTACACTCTCACCAGCTCAG TGACTGTACCCTCCAGCACCTGGTCCAGCCAGGCCGTCACCTGCAACGTAGCCCACCCGGCCAGCAGCACCAAGGTGGACAA GAAAATTGGTGAGAGAACAACCAGGGGATGAGGGGCTCACTAGAGGTGAGGATAAGGCATTAGATTGCCTACACCAACCAGGG TGGGCAGACATCACCAGGGAGGGGGCCTCAGCCCAGGAGACCAAAAATTCTCCTTTGTCTCCCTTCTGGAGATTTCTATGTCCT TTACACCCATTTATTAATATTCTGGGTAAGATGCCCTTGCATCATGACATACAGAGGCAGACTAGAGTATCAACCTGCAAAAGGTC ATACCCAGGAAGAGCCTGCCATGATCCCACACCAGAACCAACCTGGGGCCTTCTCACCTATAGACCATACTAACACACAGCCTT CTCTCTGCAGTGCCAAG |
| Sortag Histag. Stop | GGAAGGAGGCGGAGGCAGCCTGCCGGAAACCGGCGGCCATCATCATCATCATCATTGA |
| IRES | GGATCCCAATTGCTCGAGGCCCCTCTCCCTCCCCCCCCCCTAACGTTACTGGCCGAAGCCGCTTGGAATAAGGCCGGTGTGCG TTTGTCTATATGTTATTTTCCACCATATTGCCGTCTTTTGGCAATGTGAGGGCCCGGAAACCTGGCCCTGTCTTCTTGACGAGCAT TCCTAGGGGTCTTTCCCCTCTCGCCAAAGGAATGCAAGGTCTGTTGAATGTCGTGAAGGAAGCAGTTCCTCTGGAAGCTTCTTG AAGACAAACAACGTCTGTAGCGACCCTTTGCAGGCAGCGGAACCCCCCACCTGGCGACAGGTGCCTCTGCGGCCAAAAGCCAC GTGTATAAGATACACCTGCAAAGGCGGCACAACCCCAGTGCCACGTTGTGAGTTGGATAGTTGTGGAAAGAGTCAAATGGCTCT CCTCAAGCGTATTCAACAAGGGGCTGAAGGATGCCCAGAAGGTACCCCATTGTATGGGATCTGATCTGGGGCCTCGGTGCACA TGCTTTACATGTGTTTAGTCGAGGTTAAAAAAACGTCTAGGCCCCCCGAACCACGGGGACGTGGTTTTCCTTTGAAAAACACGAT GATAATATGGCCACAGAATTCGCCACC |
| *Bsr* | TGGCCAAGCCTTTGTCTCAAGAAGAATCCACCCTCATTGAAAGAGCAACGGCTACAATCAACAGCATCCCCATCTCTGAAGACTA CAGCGTCGCCAGCGCAGCTCTCTCTAGCGACGGCCGCATCTTCACTGGTGTCAATGTATATCATTTTACTGGGGGACCTTGTGC AGAACTCGTGGTGCTGGGCACTGCTGCTGCTGCGGCAGCTGGCAACCTGACTTGTATCGTCGCGATCGGAAATGAGAACAGGG GCATCTTGAGCCCCTGCGGACGGTGCCGACAGGTGCTTCTCGATCTGCATCCTGGGATCAAAGCCATAGTGAAGGACAGTGAT GGACAGCCGACGGCAGTTGGGATTCGTGAATTGCTGCCCTCTGGTTATGTGTGGGAGGGCTAAGTACTAGTCGA |
| SV40 polyA terminati on | GTACTAGTCGACTGTGCCTTCTAGTTGCCAGCCATCTGTTGTTTGCCCCTCCCCCGTGCCTTCCTTGACCCTGGAAGGTGCCAC TCCCACTGTCCTTTCCTAATAAAATGAGGAAATTGCATCGCATTGTCTGAGTAGGTGTCATTCTATTCTGGGGGGTGGGGTGGGG CAGGACAGCAAGGGGGAGGATTGGGAAGACAATAGCAGGCATGCTGGGGATGCGGTGGGCTCTATGGAGATCTTGTACA |
| 3'HA | GGTAAGTCACTAGGACTATTACTCCAGCCCCAGATTCAAAAAATATCCTCAGAGGCCCATGTTAGAGGATGACACAGCTATTGAC CTATTTCTACCTTTCTTCTTCATCTACAGGCTCAGAAGTATCATCTGTCTTCATCTTCCCCCCAAAGACCAAAGATGTGCTCACCAT CACTCTGACTCCTAAGGTCACGTGTGTTGTGGTAGACATTAGCCAGAATGATCCCGAGGTCCGGTTCAGCTGGTTTATAGATGA CGTGGAAGTCCACACAGCTCAGACTCATGCCCCGGAGAAGCAGTCCAACAGCACTTTACGCTCAGTCAGTGAACTCCCCATCGT GCACCGGGACTGGCTCAATGGCAAGACGTTCAAATGCAAAGTCAACAGTGGAGCATTCCCTGCCCCCATCGAGAAAAGCATCTC CAAACCCGAAGGTGGGAGCAGCAGGGTGTGTGGTGTAGAAGCTGCAGTAGGCCATAGACAGAGCTTGACTTAACTAGACTT |

**Table S2. Isotype donor constructs for HDR.** Table displays sequences of each feature of donor constructs used to change the isotype of rat IgG2a hybridomas. The 5' HA, IRES-*Bsr*-PolyA and 3' HA are constant in each HDR construct, while the isotype features vary between constructs. The native splice acceptor which is removed is indicated in 3' HA.

| Isotype | Sequence |
|---|---|
| 5' HA | AGAAAGATCTGAGTAGAACCAAGGTAAAAAGTGTGGGTAAAAACACATGTTCACAGGCCTGGCTGACATGATGCTGGGCACGTATGGAGGCAAAGTCAAGAGGGCAGTG<br>TAAGGGCCAGAAGTGAATCCTGACCCAAGAATAGAGAGTGCTAAACCTACGTAGATCGAAGCCAACTAAAAAGACAAGCTACAAAACGAAGCTAAGGCCAGAGATCTTGG<br>ACTGTGAAGAGTTCAGAGAACCTAGGATCAGGAACCATTAGTAACAGGCCAAGGAAGATAGAAGCTGCCTAGGACTTGGCAAGAGCCAACATGGTTGGACTGGAAAAGAA<br>AGGAGGAGACAGAAGACAGGAGAGATGTGCCAACTTGATTTTGGGCTTCACTGTTGTCCATACTGTGTGCAGCCATATGGCCCACAGATAACAGGTTTAGCCGAGGAACA<br>CAGATACCCACATTGGACAATGGTGGGGGAACACAGATACCCATACTACAGGGCTCTTTAGGGCATTTCCTGAAAGTGTACTAGGAGTGGGACTGGGCTCAAAGGGATTA<br>GGTGTGATCTGGCCTGGTGAGGCTGACATTGGCAAGCCCAATGGTTGGGTGTTGCCTCCTCCATGT |
| mIgG1 | GCTAGCGATCGCAGGCGCAATCTTCGCATTTCTTTTTTCCAGCAAAGACCACACCACCTTCTGTGTACCCACTCGCACCAGGCAGCGCCGCTCAAACCAACAGTATGGTG<br>ACCTTGGGGTGTCTTGTGAAGGGCTACTTTCCCGAGCCCGTTACCGTCACCTGGAACTCCGGGTCTCTCTCAAGCGGCGTTCACACCTTCCCCGCCGTACTGCAATCAGA<br>CCTCTATACCCTGTCTTCCAGCGTCACTGTACCCAGTTCCACCTGGCCCAGTGAAACTGTCACATGTAATGTCGCACACCCAGCTTCCTCCACCAAAGTGGACAAAAAGAT<br>CGTGCCTAGGGACTGTGGATGCAAGCCATGTATCTGCACTGTCCCCGAGGTTTCAAGTGTATTTATCTTCCCCCCAAAGCCCAAAGATGTCCTCACCATAACACTCACACC<br>CAAGGTGACCTGCGTGGTGGTCGATATCAGTAAAGATGACCCCGAGGTCCAATTCAGCTGGTTTGTGGATGATGTAGAGGTCCACACTGCTCAGACTCAGCCACGGGAA<br>GAGCAGTTTAACTCAACATTTCGGAGTGTGTCCGAACTGCCAATCATGCACCAAGATTGGCTCAATGGGAAAGAATTCAAATGCAGGGTGAATAGTGCCGCCTTTCCTGCA<br>CCAATAGAGAAAACCATATCAAAGACAAAGGGCAGGCCCAAGGCCCCTCAAGTCTATACTATACCTCCACCCAAGGAGCAAATGGCTAAAGATAAGGTAAGCCTCACCTG<br>CATGATCACAGATTTTTTCCCAGAAGACATAACTGTCGAATGGCAATGGAACGGCCAACCTGCAGAAAATTACAAGAATACTCAGCCTATTATGGACACCGATGGCAGCTA<br>CTTCGTGTATTCAAAACTGAACGTACAGAAATCTAATTGGGAAGCAGGGAATACATTCACATGCTCTGTGTTGCATGAAGGGCTCCATAACCACCACACTGAAAAGAGCTT<br>GAGCCACTCTCCCGGAAAGCGTACGGGAGGCGGAGGCAGCCTGCCGGAAACCGGCGGCCATCATCATCATCATCATTGA |
| mIgG2a | GCTAGCGATCGCAGGCGCAATCTTCGCATTTCTTTTTTCCAGCTAAGACTACCGCTCCTAGCGTTTATCCTTTGGCCCCTGTATGTGGGGACACCACTGGCTCTTCTGTAA<br>CTCTGGGCTGTTTGGTGAAAGGGTACTTTCCAGAACCTGTAACTTTGACATGGAATTCAGGGTCTCTCTCTTCAGGAGTACATACTTTTCCCGCAGTTTTGCAAAGCGATCT<br>TTACACACTTAGTTCCTCCGTCACCGTTACTTCTTCTACCTGGCCCAGTCAGTCCATAACATGTAACGTGGCCCATCCTGCCAGCAGCACAAAAGTAGACAAAAAAATCGAA<br>CCTAGAGGCCCTACTATTAAGCCCTGCCCACCATGCAAATGCCCAGCCCCAAATCTCCTCGGTGGTCCTAGCGTCTTCATCTTCCCCCCCAAGATTAAGGATGTGCTGAT<br>GATTTCATTGAGCCCAATTGTCACATGTGTGGTCGTGGATGTGTCAGAGGATGACCCTGACGTGCAAATATCTTGGTTTGTAAATAACGTAGAGGTGCATACCGCTCAGAC<br>TCAGACTCACCGGGAGGACTATAATAGCACTCTCAGGGTGGTCTCCGCACTTCCAATTCAGCACCAGGACTGGATGTCCGGCAAAGAGTTCAAGTGTAAAGTCAATAACA<br>AGGATTTGCCCGCACCAATAGAACGGACCATCTCTAAACCTAAAGGGAGTGTACGCGCCCCACAGGTTTACGTGCTGCCCCCACCCGAGGAGGAAATGACCAAAAAGCA<br>GGTGACACTCACCTGCATGGTTACCGATTTTATGCCCGAAGACATATATGTTGAGTGGACTAACAACGGGAAGACCGAGCTGAATTATAAAAATACCGAACCCGTTTTGGA<br>CTCAGATGGCTCATACTTCATGTACTCCAAACTCCGGGTAGAGAAAAAGAACTGGGTTGAAAGAAACAGCTACTCATGCAGCGTGGTGCATGAGGGGCTCCACAATCATC<br>ATACCACCAAGTCTTTCTCACGGACACCTGGGAAACGTACGGGAGGCGGAGGCAGCCTGCCGGAAACCGGCGGCCATCATCATCATCATCATTGA |
| mIgG2b | GCTAGCGATCGCAGGCGCAATCTTCGCATTTCTTTTTTCCAGCTAAAACAACACCCCCATCAGTCTATCCACTGGCCCCTGGGTGTGGAGATACAACTGGTTCCTCTGTGA<br>CTCTGGGATGCCTGGTCAAGGGCTACTTCCCTGAGTCAGTGACTGTGACTTGGAACTCTGGATCCCTGTCCAGCAGTGTGCACACCTTCCCAGCTCTCCTGCAGTCTGGA<br>CTCTACACTATGAGCAGCTCAGTGACTGTCCCCTCCAGCACCTGGCCAAGTCAGACCGTCACCTGCAGCGTTGCTCACCCAGCCAGCAGCACCACGGTGGACAAAAAAC<br>TTGAGCCCAGCGGGCCCATTTCAACAATCAACCCCTGTCCTCCATGCAAGGAGTGTCACAAATGCCCAGCTCCTAACCTGGAGGGTGGACCATCCGTCTTCATCTTCCCT<br>CCAAATATCAAGGATGTACTCATGATCTCCCTGACACCCAAGGTCACGTGTGTGGTGGTGGATGTGAGCGAGGATGACCCAGACGTCCGGATCAGCTGGTTTGTGAACAA<br>CGTGGAAGTACACACAGCTCAGACACAAACCCATAGAGAGGATTACAACAGTACTATCCGGGTGGTCAGTGCCCTCCCCATCCAGCACCAGGACTGGATGAGTGGCAAG<br>GAGTTCAAATGCAAGGTCAACAACAAAGACCTCCCATCACCCATCGAGAGAACCATCTCAAAAATTAAAGGGCTAGTCAGAGCTCCACAAGTATACATCTTGCCGCCACCA<br>GCAGAGCAGTTGTCCAGGAAAGATGTCAGTCTCACTTGCCTGGTCGTGGGCTTCAACCCTGGAGACATCAGTGTGGAGTGGACCAGCAATGGGCATACTGAGGAGAACT<br>ACAAGGACACCGCACCAGTCCTGGACTCTGACGGTTCTTACTTCATATACAGCAAGCTCGATATAAAAACAAGCAAGTGGGAGAAAACAGATTCCTTCTCATGCAACGTGA<br>GACACGAGGGTCTGAAAAATTACTACCTGAAGAAGACCATCTCCCGGTCTCCGGGTAAACGTACGGGAGGCGGAGGCAGCCTGCCGGAAACCGGCGGCCATCATCATC<br>ATCATCATTGA |
| mIgG3 | GCTAGCGATCGCAGGCGCAATCTTCGCATTTCTTTTTTCCAGCTACAACAACAGCCCCATCTGTCTATCCCTTGGTCCCTGGCTGCAGTGACACATCTGGATCCTCGGTGA<br>CACTGGGATGCCTTGTCAAAGGCTACTTCCCTGAGCCGGTAACTGTAAAATGGAACTATGGAGCCCTGTCCAGCGGTGTGCGCACAGTCTCATCTGTCCTGCAGTCTGGG<br>TTCTATTCCCTCAGCAGCTTGGTGACTGTACCCTCCAGCACCTGGCCCAGCCAGACTGTCATCTGCAACGTAGCCCACCCAGCCAGCAAGACTGAGTTGATCAAGAGAAT<br>CGAGCCTAGAATACCCAAGCCCAGTACCCCCCCAGGTTCTTCATGCCCACCTGGTAACATCTTGGGTGGACCATCCGTCTTCATCTTCCCCCCAAAGCCCAAGGATGCAC<br>TCATGATCTCCCTAACCCCCAAGGTTACGTGTGTGGTGGTGGATGTGAGCGAGGATGACCCAGATGTCCATGTCAGCTGGTTTGTGGACAACAAAGAAGTACACACAGCC<br>TGGACACAGCCCCGTGAAGCTCAGTACAACAGTACCTTCCGAGTGGTCAGTGCCCTCCCCATCCAGCACCAGGACTGGATGAGGGGCAAGGAGTTCAAATGCAAGGTCA<br>ACAACAAAGCCCTCCCAGCCCCCATCGAGAGAACCATCTCAAAACCCAAAGGAAGAGCCCAGACACCTCAAGTATACACCATACCCCCACCCCGTGAACAAATGTCCAAG<br>AAGAAGGTTAGTCTGACCTGCCTGGTCACCAACTTCTTCTCTGAAGCCATCAGTGTGGAGTGGGAAAGGAACGGAGAACTGGAGCAGGATTACAAGAACACTCCACCCAT<br>CCTGGACTCAGATGGGACCTACTTCCTCTACAGCAAGCTCACTGTGGATACAGACAGTTGGTTGCAAGGAGAAATTTTTACCTGCTCCGTGGTGCATGAGGCTCTCCATAA<br>CCACCACACACAGAAGAACCTGTCTCGCTCCCCTGGTAAACGTACGGGAGGCGGAGGCAGCCTGCCGGAAACCGGCGGCCATCATCATCATCATCATTGA |
| mIgA | GCTAGCGATCGCAGGCGCAATCTTCGCATTTCTTTTTTCCAGGCTCAGAGTCTGCGAGAAATCCCACCATCTACCCACTGACACTCCCACCAGCTCTGTCAAGTGACCCAG<br>TGATAATCGGCTGCCTGATTCACGATTACTTCCCTTCCGGCACGATGAATGTGACCTGGGGAAAGAGTGGGAAGGATATAACCACCGTAAACTTCCCACCTGCCCTGGCC<br>TCTGGGGGACGGTACACCATGAGCAGCCAGTTGACCCTGCCAGCTGTCGAGTGCCCAGAAGGAGAATCCGTGAAATGTTCCGTGCAACATGACTCTAACCCCGTCCAAG<br>AATTGGATGTGAATTGCTCTGGTCCTACTCCTCCTCCTCCTATTACTATTCCTTCCTGCCAGCCCAGCCTGTCACTGCAGCGGCCAGCTCTTGAGGACCTGCTCCTGGGTT<br>CAGATGCCAGCATCACATGTACTCTGAATGGCCTGAGAAATCCTGAGGGAGCTGTCTTCACCTGGGAGCCCTCCACTGGGAAGGATGCAGTGCAGAAGAAAGCTGTGCA<br>GAATTCCTGCGGCTGCTACAGTGTGTCCAGCGTCCTGCCTGGCTGTGCTGAGCGCTGGAACAGTGGCGCATCATTCAAGTGCACAGTTACCCATCCTGAGTCTGGCACC<br>TTAACTGGCACAATTGCCAAAGTCACAGTGAACACCTTCCCACCCCAGGTCCACCTGCTACCGCCGCCGTCGGAGGAGCTGGCCCTGAATGAGCTCTTGTCCCTGACATG<br>CCTGGTGCGAGCTTTCAACCCTAAAGAAGTGCTGGTGCGATGGCTGCATGGAAATGAGGAGCTGTCCCCAGAAAGCTACCTAGTGTTTGAGCCCCTAAAGGAGCCAGGC<br>GAGGGAGCCACCACCTACCTGGTGACAAGCGTGTTGCGTGTATCAGCTGAAACCTGGAAACAGGGTGACCAGTACTCCTGCATGGTGGGCCACGAGGCCTTGCCCATGA<br>ACTTCACCCAGAAGACCATCGACCGTCTGTCGGGTAAACCCACCAATGTCAGCGTGTCTGTGATCATGTCAGAGGGAGATGGCATCTGCTACCGTACGGGAGGCGGAGG<br>CAGCCTGCCGGAAACCGGCGGCCATCATCATCATCATCATTGA |
| mIgG2a silent | GCTAGCGATCGCAGGCGCAATCTTCGCATTTCTTTTTTCCAGCTAAGACTACCGCTCCTAGCGTTTATCCTTTGGCCCCTGTATGTGGGGACACCACTGGCTCTTCTGTAA<br>CTCTGGGCTGTTTGGTGAAAGGGTACTTTCCAGAACCTGTAACTTTGACATGGAATTCAGGGTCTCTCTCTTCAGGAGTACATACTTTTCCCGCAGTTTTGCAAAGCGATCT<br>TTACACACTTAGTTCCTCCGTCACCGTTACTTCTTCTACCTGGCCCAGTCAGTCCATAACATGTAACGTGGCCCATCCTGCCAGCAGCACAAAAGTAGACAAAAAAATCGAA<br>CCTAGAGGCCCTACTATTAAGCCCTGCCCACCATGCAAATGCCCAGCCCCAAATgcCgcCGGTGGTCCTAGCGTCTTCATCTTCCCCCCCAAGATTAAGGATGTGCTGATG<br>ATTTCATTGAGCCCAATTGTCACATGTGTGGTCGTGGATGTGTCAGAGGATGACCCTGACGTGCAAATATCTTGGTTTGTAAATAACGTAGAGGTGCATACCGCTCAGACT<br>CAGACTCACCGGGAGGACTATgccAGCACTCTCAGGGTGGTCTCCGCACTTCCAATTCAGCACCAGGACTGGATGTCCGGCAAAGAGTTCAAGTGTAAAGTCAATAACAAG<br>GATTTGCCCGCACCAATAGAACGGACCATCTCTAAACCTAAAGGGAGTGTACGCGCCCCACAGGTTTACGTGCTGCCCCCACCCGAGGAGGAAATGACCAAAAAGCAGG<br>TGACACTCACCTGCATGGTTACCGATTTTATGCCCGAAGACATATATGTTGAGTGGACTAACAACGGGAAGACCGAGCTGAATTATAAAAATACCGAACCCGTTTTGGACT<br>CAGATGGCTCATACTTCATGTACTCCAAACTCCGGGTAGAGAAAAAGAACTGGGTTGAAAGAAACAGCTACTCATGCAGCGTGGTGCATGAGGGGCTCCACAATCATCAT<br>ACCACCAAGTCTTTCTCACGGACACCTGGGAAACGTACGGGAGGCGGAGGCAGCCTGCCGGAAACCGGCGGCCATCATCATCATCAcCATTGA |
| IRES *Bsr* polyA | GTCGACGTCGAGGCCCCTCTCCCTCCCCCCCCCCTAACGTTACTGGCCGAAGCCGCTTGGAATAAGGCCGGTGTGCGTTTGTCTATATGTTATTTTCCACCATATTGCCG<br>TCTTTTGGCAATGTGAGGGCCCGGAAACCTGGCCCTGTCTTCTTGACGAGCATTCCTAGGGGTCTTTCCCCTCTCGCCAAAGGAATGCAAGGTCTGTTGAATGTCGTGAA<br>GGAAGCAGTTCCTCTGGAAGCTTCTTGAAGACAAACAACGTCTGTAGCGACCCTTTGCAGGCAGCGGAACCCCCCACCTGGCGACAGGTGCCTCTGCGGCCAAAAGCCA<br>CGTGTATAAGATACACCTGCAAAGGCGGCACAACCCCAGTGCCACGTTGTGAGTTGGATAGTTGTGGAAAGAGTCAAATGGCTCTCCTCAAGCGTATTCAACAAGGGGCT<br>GAAGGATGCCCAGAAGGTACCCCATTGTATGGGATCTGATCTGGGGCCTCGGTGCACATGCTTTACATGTGTTTAGTCGAGGTTAAAAAAACGTCTAGGCCCCCCGAACC<br>ACGGGGACGTGGTTTTCCTTTGAAAAACACGATGATAATATGGCCACAGAATTCGCCACCATGGCCAAGCCTTTGTCTCAAGAAGAATCCACCCTCATTGAAAGAGCAACG<br>GCTACAATCAACAGCATCCCCATCTCTGAAGACTACAGCGTCGCCAGCGCAGCTCTCTCTAGCGACGGCCGCATCTTCACTGGTGTCAATGTATATCATTTTACTGGGGGA<br>CCTTGTGCAGAACTCGTGGTGCTGGGCACTGCTGCTGCTGCGGCAGCTGGCAACCTGACTTGTATCGTCGCGATCGGAAATGAGAACAGGGGCATCTTGAGCCCCTGCG<br>GACGGTGCCGACAGGTGCTTCTCGATCTGCATCCTGGGATCAAAGCCATAGTGAAGGACAGTGATGGACAGCCGACGGCAGTTGGGATTCGTGAATTGCTGCCCTCTGG<br>TTATGTGTGGGAGGGCTAAGTACTAGTCGAGTGTGCCTTCTAGTTGCCAGCCATCTGTTGTTTGCCCCTCCCCCGTGCCTTCCTTGACCCTGGAAGGTGCCACTCCCACT<br>GTCCTTTCCTAATAAAATGAGGAAATTGCATCGCATTGTCTGAGTAGGTGTCATTCTATTCTGGGGGGTGGGGTGGGGCAGGACAGCAAGGGGGAGGATTGGGAAGACA<br>ATAGCAGGCATGCTGGGGATGCGGTGGGCTCTATGGAGATCTTGT |
| 3' HA | TGTACAACTTGGGGAGGGTACAAAATGGAGGACTTGTAGGAGCTTGGGTCCAGACCTGTCAGACAAAATGATCACGCATACTTATTCTTGT***AGC***TGAAACAACAGCCCCAT<br>CTGTCTATCCACTGGCTCCTGGAACTGCTCTCAAAAGTAACTCCATGGTGACCCTGGGATGCCTGGTCAAGGGCTATTTCCCTGAGCCAGTCACCGTGACCTGGAACTCT<br>GGAGCCCTGTCCAGCGGTGTGCACACCTTCCCAGCTGTCCTGCAGTCTGGACTCTACACTCTCACCAGCTCAGTGACTGTACCCTCCAGCACCTGGTCCAGCCAGGCCG<br>TCACCTGCAACGTAGCCCACCCGGCCAGCAGCACCAAGGTGGACAAGAAAATTGGTGAGAGAACAACCAGGGGATGAGGGGCTCACTAGAGGTGAGGATAAGGCATTA<br>GATTGCCTACACCAACCAGGGTGGGCAGACATCACCAGGGAGGGGGCCTCAGCCCAGGAGACCAAAAATTCTCCTTTGTCTCCCTTCTGGAGATTTCTATGTCCTTTACA<br>CCCATTTATTAATATTCT |

**Table S3. FcγR affinity values of MIH5 Fc variants and comparison to literature.** Affinity quantification by $K_D$ (μM) of CRISPR/HDR antibodies for mFcγRI, FcγRIIb and mFcγRIV (bold) comparison to described values in literature.

| Fc variant | mFcγRI | mFcγRIIb | mFcγRIV |
|---|---|---|---|
| **mIgG1** | **-/-**<br>-/- [S1-S6] | **0.26 ± 0.05**<br>0.15[S1]; 0.30[S2] ; 0.83[S3]; 0.17[S4]; 0.25[S5] | **-/-**<br>-/- [S1-S6] |
| **mIgG2a** | **0.017 ± 0.007**<br>0.012 [S1]; 0.006 [S2]; 0.026 [S3]; 0.018 [S4]; 0.033 [S6]; 0.013 - 0.022 [S7] | **0.66 ± 0.20**<br>0.69[S1]; 2.4[S2]; 1.8[S3] | **0.13 ± 0.02**<br>0.060[S1]; 0.035[S2]; 0.071[S3]; 0.010[S4] |
| **mIgG2b** | **-/-**<br>-/- [S1, S3, S7]; 10[C]; 0.021[S7] | **1.24 ± 0.06**<br>0.83[S1]; 0.45[S2]; 0.91[S3]; 0.26[S5] | **0.20 ± 0.02**<br>0.12[S1]; 0.059[S2]; 0.063[S3]; 0.034[S5] |
| **mIgA** | **-/-**<br>-/-[S6] | **-/-**<br>-/-[S6] | **-/-**<br>-/-[S6] |
| **mIgG2a$_{silent}$** | **-/-**<br>-/-[S8] | **-/-**<br>-/-[S8] | **-/-**<br>-/-[S8] |

References;


**S1.** Dekkers, G. et al. Affinity of human IgG subclasses to mouse Fc gamma receptors. MAbs 9, 767–773 (2017).

**S2**. Nimmerjahn, F., Bruhns, P., Horiuchi, K. & Ravetch, J. V. FcγRIV: A Novel FcR with Distinct IgG Subclass Specificity. Immunity 23, 41–51 (2005).

**S3.** Baudino, L. et al. Impact of a three amino acid deletion in the CH2 domain of murine IgG1 on Fc-associated effector functions. J. Immunol. 181, 4107–12 (2008).

**S4.** White, A. L. et al. Interaction with FcγRIIB is critical for the agonistic activity of anti-CD40 monoclonal antibody. J. Immunol. 187, 1754–63 (2011).

**S5.** Kaneko, Y., Nimmerjahn, F. & Ravetch, J. V. Anti-Inflammatory Activity of Immunoglobulin G Resulting from Fc Sialylation. Science (80-. ). 313, 670–673 (2006).

**S6.** Bruhns, P. & Jönsson, F. Mouse and human FcR effector functions. Immunol. Rev. 268, 25–51 (2015).

**S7.** Gavin, A. L., Leiter, E. H. & Hogarth, P. M. Mouse FcgammaRI: identification and functional characterization of five new alleles. Immunogenetics 51, 206–11 (2000). **S8.** Arduin, E. et al. Highly reduced binding to high and low affinity mouse Fc gamma receptors by L234A/L235A and N297A Fc mutations engineered into mouse IgG2a. Mol. Immunol. 63, 456–463 (2015).
